# Microfluidics-free single-cell genomics reveals complex central-peripheral immune crosstalk in the mouse brain during peripheral inflammation

**DOI:** 10.1101/2023.10.05.561054

**Authors:** Jake Sondag Boles, Oihane Uriarte Huarte, Malú Gámez Tansey

## Abstract

Inflammation is a realized detriment to brain health in a growing number of neurological diseases, but querying neuroinflammation in its cellular complexity remains a challenge. This manuscript aims to provide a reliable and accessible strategy for examining the brain’s immune system. We compare the efficacy of cell isolation methods in producing ample and pure immune samples from mouse brains. Then, with the high-input single-cell genomics platform PIPseq, we generate a rich neuroimmune dataset containing microglia and many peripheral immune populations. To demonstrate this strategy’s utility, we interrogate the well-established model of LPS-induced neuroinflammation with single-cell resolution. We demonstrate the activation of crosstalk between microglia and peripheral phagocytes and highlight the unique contributions of microglia and peripheral immune cells to neuroinflammation. Our approach enables the high-depth evaluation of inflammation in longstanding rodent models of neurological disease to reveal novel insight into the contributions of the immune system to brain health.

## Introduction

Inflammation is becoming a key component in a growing number of neuropsychiatric and neurodegenerative diseases^1–3^, and its role in brain diseases ranges from protective to injurious. In the brain, microglia, as the resident phagocytes, are often the first to be implicated in pathogenic neuroinflammation^4^. However, the infiltration of peripheral innate and adaptive immune cells has been observed in human diseases as well as rodent models created to study those diseases^5^. Moreover, recent single-cell omics studies emphasize the heterogeneity that can exist in a single cell type^6,7^. These critical observations compel a re-examination of neurological diseases and their pre-clinical models with finer resolution and an appreciation for resident and non-resident immune cells.

Flow cytometry has been the gold standard method used to study immune cells because it enables the identification of specific cell types and their functional states^8^. However, this technique has an inherent bias due to the limited number of channels of a flow cytometer, requiring the user to prioritize targets. The heterogeneity of the immune cells may limit the utility of selected targets, as activation states themselves may disrupt protein marker presentation^7,9^. Additionally, detecting rarer cell populations with flow cytometry is difficult due to technical noise^10^, resulting in the user missing smaller but non-trivial components of the brain microenvironment. Mass cytometry enables higher parameterization and circumvents some of the limitations of flow cytometry, but technical barriers limit the widespread use of this technique^11,12^.

Single-cell genomics is the current state-of-the-art technique, permitting the study of cell-specific transcriptomic profiles and inferred communication between different populations within the sample of single cells. In addition, the unbiased nature of this approach enables the identification of novel-cell subtypes or states, thereby adding greater value to the examination of immune phenotypes using single-cell approaches^13^. However, this methodology has its own limitations. Assaying immune cells from the brain remains difficult as cell isolation strategies vary in their efficiency (yield and viability). Furthermore, fluorescence-activated cell sorting (FACS) of CD45+ cells is generally employed to enhance the purity of samples^14,15^, a step which exhibits the inherent limitations of flow cytometry^12^ and may be difficult to access depending on institutional infrastructure. To combat these limitations, here we compare two different immune cell-isolation strategies and explore the effectiveness of magnetic separation (MS), which can be performed at the bench with minimal equipment in lieu of FACS and yields pure immune samples from mouse brain. Because high reagent and equipment cost can be prohibitive, we employed the novel microfluidics-free platform that uses particle-templated instant partitions (PIPseq) to lessen the cost of single-cell genomics^16^. An added advantage to the PIPseq platform is its scalability and high cell-input capacity^16,17^, which bolsters our ability to capture and examine immune cells present in low numbers in or surrounding the brain.

To showcase the effectiveness of our accessible approach combining MS and PIPseq, we sought to interrogate the brain’s immune system during peripheral lipopolysaccharide (LPS)-induced neuroinflammation. Using this technique, we replicated key immunological observations previously reported in this model of peripheral inflammation, including the infiltration of peripheral myeloid cells into the brain^18,19^. We infer key signaling patterns between microglia and these infiltrating phagocytes that likely underlie extravasation, suggesting that LPS-exposed microglia preferentially activate communication with non-resident immune cells. Finally, we showcase the unique and overlapping transcriptomic programs of brain immune cells during LPS-induced neuroinflammation, characterized by the cell-specific regulation of many disease-related genes. This novel strategy will enable the investigation of immune crosstalk in animal models of neurological disease across the neurodegeneration field, enabling the rapid discovery of novel targets and signaling pathways related to inflammation that may be fruitful therapeutic targets to rescue vulnerable neuronal populations and prevent or mitigate brain injury.

## Methods

### Mice

C57Bl/6J (Strain #000664) and progranulin (*Grn*-/-) deficient mice (B6(Cg)-*Grn^tm^*^1^*^.1Aidi^*/J, Strain #013175) were obtained from Jackson Laboratories and housed in individually ventilated cages and maintained with *ad libitum* access to standard rodent chow on a 12:12 light-dark cycle in a conventional animal facility in the McKnight Brain Institute vivarium at the University of Florida. All procedures were approved by the University of Florida Institutional Animal Care and Use Committee and followed the *Guide for the Care and Use of Laboratory Animals* from the National Institutes of Health (NIH Publications No. 80-23, revised 1996).

To compare immune cell isolation procedures from brain, 4- to 10-month-old C57Bl/6J and *Grn*-/- mice were euthanized via rapid decapitation and brains were rapidly extracted. To test different concentrations of CD45 MicroBeads and CD45 fluorescent antibody clones, 5- to 10-month-old C57Bl/6J mice were euthanized via rapid decapitation and brains were rapidly extracted. Finally, to examine a widely used model of neuroinflammation with our optimized brain immune cell isolation and the PIPseq single-cell genomics platform, 3- to 5-month-old male C57Bl/6J received a single intraperitoneal (i.p.) injection of lipopolysaccharide (1.5 x 10^7^ EU/kg) from *Escherichia coli* 0111:B4 (Sigma, #L2630) or an equivalent volume of sterile saline. Twenty-four hours post-injection, mice were euthanized via rapid decapitation and brains were quickly extracted.

### Brain dissociation protocols

#### Collagenase VIII, DNase I, and Percoll

Brains were extracted from C57Bl/6J or *Grn*-/- mice and bisected. One hemisphere was finely minced, digested with 1.4U/mL collagenase VIII (Sigma, #C2139) and 1mg/mL DNase I (Sigma, #DN25) in RPMI 1640 medium (ThermoFisher, #11875119) for 15min at 37°C after which the enzymes were inactivated with the addition of 10% fetal bovine serum (FBS) in RPMI 1640 medium. Digested tissue was manually triturated using fire-polished Pasteur pipettes and filtered through a 70µm nylon cell-strainer, leaving a single-cell suspension. Tissue lysates were suspended in a solution made of 37% Percoll (Sigma, #P1644) in Hank’s balanced salt solution (HBSS-/-; ThermoFisher, #14175103). A layer of 70% Percoll in HBSS-/- was carefully deposited underneath the tissue layer and a layer of 30% Percoll in HBSS-/- was slowly dispensed on top of these two phases. This layered solution was centrifuged for 30 minutes at 400 x *g* with braking disabled to avoid collapsing the layers, after which the top phase containing cellular debris and myelin was discarded. Cells were aspirated from the lower interphase, washed in phosphate buffered saline (PBS), and carried forward to surface staining and flow cytometry.

#### Adult Brain Dissociation Kit and CD45 magnetic separation

The other hemisphere from the above mice was treated with Miltenyi Biotec Adult Brain Dissociation Kit (ABDK, #130-107-677). Hemispheres were cut into 8-12 small pieces and placed into gentleMACS C-tubes (Miltenyi Biotec, #130-093-237) with dissociation solution prepared as instructed by the manufacturer’s protocol. When cells were being isolated for single-cell RNA sequencing, 5µg/mL actinomycin D, 10µM triptolide, and 27.1µg/mL anisomycin or an equal volume of dimethyl sulfoxide (DMSO) as used previously ^20^ were added to the dissociation solution to enable the evaluation of transcriptional artifacts induced by this method in this sample type. Tissue was subjected to the 37C_ABDK_01 protocol on the gentleMACS Octo Dissociator with heaters (Miltenyi Biotec, #130-096-427). Lysate was filtered through a 70µm nyclon cell-strainer with Dulbecco’s PBS with calcium, magnesium, glucose, and pyruvate (D-PBS). Myelin and cell debris was removed using Debris Removal Solution according to manufacturer’s instructions, which involved resuspending cells in a solution of 0.9mL Debris Removal Solution and 3.1mL D-PBS, gently layering D-PBS above this solution, and centrifuging at 3000 x *g* for 10 minutes at 4°C with slower braking. The top two phases were discarded, cells were washed gently in D-PBS, and red blood cells were lysed with Red Blood Cell Removal Solution diluted 1:10 in distilled water for 10 minutes at 4°C. Lysis was quenched with the addition of 10x volume of D-PBS with 0.5% bovine serum albumin (BSA) added. Cells were resuspended in PBS in the case of a follow-up study or carried to CD45+ cell enrichment (see below).

To enrich for immune cells, cell suspensions were subjected to magnetic separation with antibodies conjugated to magnetic beads. Cells were suspended in 90µL of a buffer made of autoMACS Rinsing Solution (Miltenyi Biotec, #130-091-022) with 0.5% BSA (BB), to which 10µL CD45 MicroBeads (Miltenyi Biotec, #130-052-301) was added. A second experiment was conducted that performed a MicroBeads dilution series, with the only adjustment to this protocol being the ratio of BB to MicroBeads here. Labeling occurred over 15 min at 4°C. Labeling was quenched with the addition of 20X volume BB. Labeled cells were added to MS columns (Miltenyi Biotec, #130-042-201) in a magnetic OctoMACS Separator (Miltenyi Biotec, #130-042-108). Columns were washed thrice with BB to elute unlabeled cells, which were collected only in the second experiment, and labeled cells were eluted in BB using the provided column plungers. Cells were washed in PBS and subjected to surface labeling and flow cytometry or cell capture and library preparation.

### Multi-color flow cytometry

To compare tissue dissociation procedures, cells were transferred to a V-bottom 96-well plate (Sigma, #CLS3896-48EA) and centrifuged at 300 x *g* for 5 minutes at 4°C. Pelleted cells were incubated with an antibody cocktail described in Table S1 with LIVE/DEAD Fixable Aqua (Invitrogen, #L34957) for 20 minutes at 4°C. Cells were washed twice in PBS and fixed with eBioscience IC Fixation Buffer (Invitrogen, #00-8222-49) for 30 minutes at 4°C. After one wash, cells were resuspended in 300µL FACS buffer (0.25mM EDTA, 0.01% NaN_3_, and 0.1% BSA in PBS) and analyzed using a BD LSRFortessa. Compensation was achieved using SPHERO Supra Rainbow beads (Spherotech, Inc., #SRCP-35-2A) by setting laser voltages to achieve fluorescent intensity measurements in each channel equal to those in previous experiments. The compensation matrix was created with the acquisition of single-stained AbC Total Antibody Compensation Beads (ThermoFisher, #A10497) for antibodies or ArC Amine Reactive Compensation Beads (ThermoFisher, #A1034) for the LIVE/DEAD dye.

To evaluate different concentrations of CD45 MicroBeads, cells were stained, washed, and fixed as described, but the antibody cocktail consisted only of LIVE/DEAD Fixable Aqua dye, CD11b-PE, and one of three CD45-APC antibodies described in Table S2. These three antibodies are distinct clones of a monoclonal CD45 antibody, which we used to evaluate whether residual CD45-cells in our enriched samples could be attributed to competition between the CD45 MicroBead antibody and the antibody used during fluorescent surface labeling. These samples were analyzed on a MACSQuant10 instrument. Compensation was conducted automatically using single-stained OneComp eBeads (Invitrogen, #01-1111-42). For both experiments, a minimum of 100,000 live cell events were captured and gating and population analysis was performed in FlowJo v.10.9.0 with the assistance of fluorescence-minus-one controls.

### Single cell RNA library preparation with PIPseq and next-generation sequencing

Single-cell RNA library preparation was performed using the PIPseq T20 kit (Fluent BioSciences) following the manufacturer’s instructions (revision 7.3). Briefly, 40,000 CD45+ cells per sample were isolated and loaded into PIPs to capture 20,000 cells. Cells were lysed and barcoded mRNA was isolated to perform cDNA synthesis and amplification. cDNA quality was checked using a Qubit dsDNA High Sensitivity assay (ThermoFisher, #Q323854) and High Sensitivity DNA Kit (Agilent, #5067-4626) on a Bioanalyzer 2100 (Agilent) before library preparation was performed. Integrity of cDNA libraries were assessed with a High Sensitivity DNA Kit on a Bioanalyzer 2100 before samples were sent for 2×150bp paired-end sequencing on an Illumina NovaSeq 6000 (Illumina) at the Interdisciplinary Center for Biotechnology Research (ICBR) core at University of Florida.

#### Data pre-processing and quality control

FASTQ files were processed through the PIPseeker pipeline v.2.0, which handles barcode and paired-end read matching and trims adapters off read 2 to enable genome alignment. Genome alignment is carried out through PIPseeker using the STAR aligner^21^ and the GRCmm39 mouse reference genome (from GENCODE). The resulting BAM file is parsed and transcripts are counted to generate a dataset containing all present gene barcodes with their unique molecular indices, which is then converted to a raw count matrix that contains the counts of each unique molecule in each cell.

This feature-barcode matrix was processed with Seurat v.4^22,23^ in R v.4.2. Seurat objects were created for each sample and merged. Mitochondrial gene abundance and cell complexity (log10 transformed number of counts divided by the log10 transformed number of unique genes) were calculated for each cell. Cells with 1) less than 5% mitochondrial genes; 2) at least 250 but less than 6500 unique genes expressed; 3) at least 800 but less than 45000 total gene counts; and 4) at least a cell complexity ratio of 0.8 were carried forward. Cutoff selection was determined through iterative selection and visual examination of relevant quality control metrics.

To handle potential batch effects due to staggered cell capture days and enable cell type identification from different conditions, the dataset was integrated using reciprocal principal component analysis (RPCA). The dataset was separated by cell capture day, and each subset was normalized using SCTransform (v.2) with ‘method = “glmGamPoi”’^24^. A set of integration anchors was computed, and RPCA was performed with this anchor set. The first 100 principal components (PCs) were computed and the first 30 were carried forward. Dimensional reduction was achieved with the uniform manifold approximation and projection (UMAP) technique with ‘RunUMAP’ and its default settings through Seurat. Nearest neighbors were computed based on the PCA and cell clusters were identified with ‘resolution = 0.2’ to allow for coarse clustering and initial cell annotation.

### Cluster annotation and data cleaning

Coarsely clustered cells were annotated using Seurat’s ‘FindAllMarkers’ function and its default settings, which uses a Wilcox test to determine differentially expressed genes in each cluster compared to all other clusters. These gene lists were interrogated using CellMarker 2.0^25^ to infer cell annotations which were confirmed with the localization of canonical marker genes. Further inspection revealed microglial markers (e.g., *P2ry12*, *Hexb*) present in non-microglia clusters. These clusters were inspected individually with re-integration and sub-clustering. In some cases, a distinct group of cells could be observed that expressed genes typical to that sub-cluster but also to microglia, so these cells were regarded as doublets and removed from the dataset. After this cleaning, the dataset was reintegrated as described above, re-clustered with a low resolution, markers were generated again, and clusters were re-annotated to ensure accurate labeling.

At this point, our dataset consisted of microglia, monocytes/macrophages, B-cells, a cluster consisting of T-cells and natural killer cells, neutrophils, and several non-immune clusters. Each of these groups was run through the chooseR pipeline^26^, which performs iterative subsampling and clustering to pick the most robust clustering strategy. The microglial dataset was randomly down-sampled to leave a dataset one-eighth in size for this pipeline due to its computational intensity. The clustering resolutions that yielded the greatest median silhouette score were selected, and markers were identified using Seurat’s ‘FindAllMarkers’ with default settings. Silhouette scores were computed for each subcluster and visualized. Cell identities were designated based on these marker lists, which were applied to the full dataset. After this process, the dataset was pruned of any doublets, reassembled, and re-integrated.

### Cell proportion analysis

To evaluate whether LPS induces a change in the cellular composition captured by our pipeline, the Speckle package^27^ was used, which transforms population proportions, fits a linear model for each cell type with the predictors of interest, and estimates *p*-values with an empirical Bayes shrinkage of variances using the Limma package^28^. We transformed the proportions in our dataset with the arcsine transformation, included only *in vivo* treatment (LPS or saline) as a predictor, and present the FDR-adjusted *p*-values for each cell type. Although groups of cell types are presented separately, all 20 populations were passed through one analysis, and the *p*-values are adjusted accordingly.

### Transcriptional artifact analysis

The annotated dataset was divided into subsets for analysis of artifactual signature induction. Due to the infrequency of certain cell types, basophils, mast cells, and neutrophils were merged into one granulocyte dataset, αβ T-cells and γδ T-cells were merged into one T-cell dataset, and plasma cells and B-cells were merged into one B-lymphocyte dataset. All non-immune cells were merged into one CD45-cell dataset. All other populations were subset individually. A cross entropy test^29^ was conducted in each dataset to determine whether the UMAP embeddings of inhibitor-treated cells and vehicle-treated cells differed significantly from one another, and the Kullback-Leibler divergences and Holm-adjusted *p-*values are shown for each comparison. Artificial activation modules were taken from differential expression analyses from pseudobulked single-cell datasets in ref.^20^ and added to the cell cluster objects with Seurat’s ‘AddModuleScorè function. Artificial gene module induction was visualized using scCustomize’s (https://cran.r-project.org/web/packages/scCustomize/index.html) wrapper functions for Seurat’s visualization tools. To separate artificially activated populations, cells were clustered and artifact module scores of each cluster were plotted using scCustomize’s ‘VlnPlot_scCustom’ function and the contribution of each experimental group to each cluster was visualized with the ‘dittoBarPlot’ function from the dittoSeq package^30^, revealing subsets of cells that displayed higher expression of the artifactual gene module and were dominated by samples that did not receive inhibitors during dissociation. Artificially activated populations were removed from the full dataset before downstream analysis as were non-immune cells.

### Intercellular communication analysis

Differences in intercellular communication between brain immune cells evoked by peripheral endotoxin were inferred and analyzed using two complementary R packages. CellChat^31^ uses mass action models and differential expression analyses to infer cell-specific signaling patterns within experimental groups of interest. The law of mass action based on the average expression of a ligand in one cell type and a receptor in another cell type is used to infer the probability of communication, and the significance of this interaction is determined by whether this probability is higher than that amongst randomly permuted cell groups.

MultiNicheNetR^32^ uses the differential state analysis as described by the muscat R package^33^, which includes cell-level mixed models and statistical tests on aggregated pseudo-bulked data. This package makes sample-level inferences on intercellular communication states, allowing us to examine both the extent and variability of the differential expression of ligand-receptor pairs. Both CellChat and MultiNicheNetR employ their own manually curated databases as the foundation of their respective communication inferences.

### Pseudo-bulked differential expression analysis

For each immune cell subset, the effect of LPS on gene expression was assessed, identifying groups of differentially expressed genes (DEGs) for each cell type. Raw gene counts from each cell in each sample were aggregated using Seurat’s ‘AggregateExpression’ to create a pseudo-bulked dataset, and immune cell subtypes were separated for DEG analysis. Differential expression was done with DESeq2^34^. A gene was designated to be differentially expressed if its absolute value of the log2 transform of the fold change with LPS exceeded 2 and its Benjamini-Hochberg adjusted *p*-value was less than 0.05.

### Gene set enrichment analysis (GSEA)

A GSEA^35^ was performed in immune cell types using the ClusterProfiler package^36^. Specifically, ranked gene lists were passed through ClusterProfiler’s ‘compareClusters’ function, where Gene Ontology knowledgebase’s^37,38^ Biological Process pathways with greater than 30 genes and fewer than 300 genes were used as the gene sets of interest. Gene ranking came from DESeq2’s effect size shrinkage procedure using the ‘apeglm’ algorithm^39^. Enrichment scores calculated during the random walk were normalized based on gene set and overlap size, and *p*-values were adjusted using the Benjamini-Hochberg correction. To compare the enrichment of biological pathways between immune cell types during LPS exposure, representative terms from the top 30 pathways with the lowest corrected *p*-value were selected from each cell type.

### AD and PD gene list curation

To examine the cell-specific expression of risk genes associated with neurodegenerative disease, gene lists were assembled from genome wide association studies (GWAS) and meta-GWAS. A set of 31 genes associated with AD was used as in ref.^40^ and ref.^41^. For PD, the 62 unique eQTL-nominated genes from ref.^42^ were sorted on the meta-*p*-value that included random effects. Both gene lists were converted from human symbols to mouse symbols, leaving 30 orthologs for AD and 59 orthologs for PD.

### Statistical analysis

All statistical analyses were performed in R v4.2-4.3. When evaluating the efficacy of different brain dissociation techniques, data were analyzed with a two-way mixed ANOVA using the Afex package (https://cran.r-project.org/web/packages/afex/index.html), with genotype as a between-subjects predictor and protocol as a within-subjects predictor. All pairwise comparisons were made using the Emmeans package (https://cran.r-project.org/web/packages/emmeans/index.html) with Tukey’s correction, and compact letter displays (CLD) were generated with the Multcomp package (https://cran.r-project.org/web/packages/multcomp/index.html). When evaluating different concentrations of CD45 MicroBeads and CD45 antibody clones, ANOVAs were run separately within each fraction (positive fraction, negative fraction, or untreated with MicroBeads). In the positive and negative fractions, two-way mixed ANOVAs were performed with antibody clone as the within-subjects factor and MicroBead dilution was the between-subjects factor. In the untreated samples, antibody clone was the single within-subjects factor in a one-way ANOVA. All pairwise comparisons were made within each fraction with Tukey’s correction and CLDs were generated. The Geisser-Greenhouse correction was used when a departure from sphericity was observed in our data according to the Performance package’s (https://cran.r-project.org/web/packages/performance/index.html) ‘check_sphericity’, which uses Mauchley’s test. All other statistical analyses performed within our single cell RNA sequencing analysis are described above. Tables displaying ANOVA and cross entropy test results were generated using the GT package (https://gt.rstudio.com/).

## Results

### Comparison of brain immune cell isolation methodologies reveals the superiority of immune cell enrichment by magnetic separation

To establish a reproducible and accessible benchtop pipeline to interrogate the brain’s immune system, we first evaluated different strategies to isolate immune cell subsets from the rodent brain without FACS. We compared a method using DNase I and collagenase VIII digestion coupled with immune cell enrichment via a Percoll gradient, a technique used in our lab and many others^43^, and dissociation with the Adult Brain Dissociation Kit (ABDK), which uses the OctoMACS for automated dissociation, followed by magnetic separation (MS) of CD45+ cells. We used C57Bl/6J mice and mice lacking progranulin (*Grn*-/-) to compare the impact of each method on cell viability, as we have previously observed lower viability of immune cells isolated from *Grn*-/- brains when using enzymatic dissociation. After cells were isolated with both methodologies, cells were immunolabelled and immune populations were quantified by flow cytometry (**Figure 1A**). The ABDK with CD45 MS captured significantly more cells in C57Bl/6J mice but not *Grn*-/- mice than collagenase VIII/DNase I with Percoll gradient isolation (Figure 1B, Table 1). As we have seen before, *Grn*-/- cells exhibit lower viability after dissociation, but this does not differ between protocols, suggesting that neither technique is ideal for sensitive cells (**Figure 1C**, **Table 1**). We investigated differences in the yield of specific populations using the gating strategy shown in **Figure S1A**. Importantly, the ABDK with CD45 MS yielded a purer immune sample, with a lower proportion of CD45-cells (**Figure 1D**, **Table 1**). We did not observe any significant changes in the raw number of T- and B-cells, CD45-cells, or neutrophils due to protocol (**Figure S1B-F, Table 1),** likely due to low counts of T-cells, B-cells, and neutrophils in the brain overall. A similar raw yield of CD45-cells between techniques coupled with the overall greater yield from the ABDK with CD45 MS underscores the earlier observation that this procedure captures a purer immune cell sample. We observed that the ABDK method increased the yield of dendritic cells, monocytes/macrophages, and microglia, but *Grn* deficiency stunted these yields (**Figure S1F-H, Table 1**). Together, these findings indicate that the ABDK dissociation method coupled with CD45 MS resulted in a higher and purer yield than collagenase/DNase dissociation with Percoll isolation.

**Figure 1:**
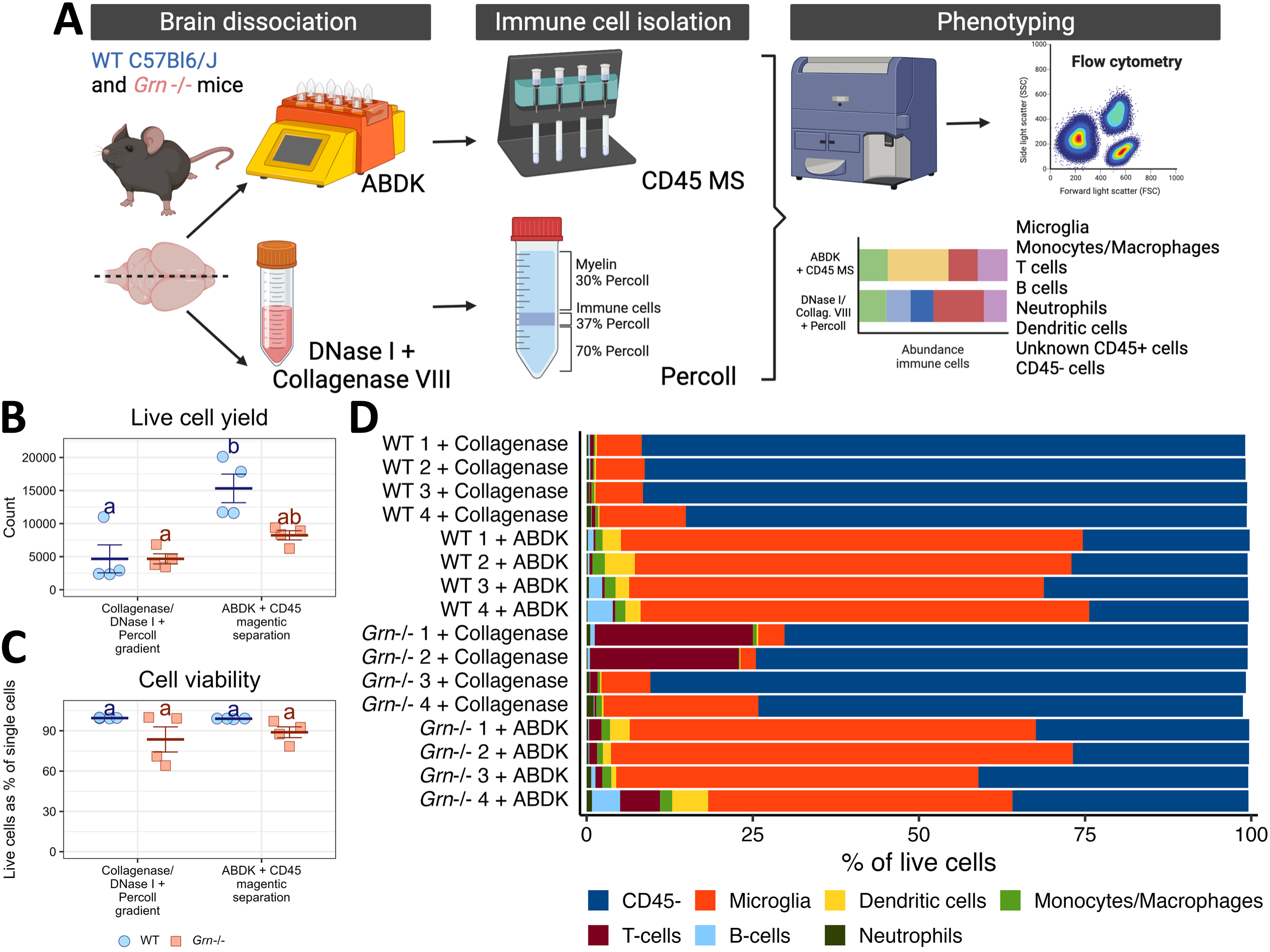
Comparison of brain immune cell isolation methodologies reveals the superiority of immune cell enrichment by magnetic separation. A) Experimental workflow for enzymatic dissociation and cell enrichment techniques. B) Live cell counts, where blue circles are B6 mice and orange squares are *Grn*-/- mice. C) Cell viability, shown as the percentage of single cells that are alive, coded the same as in (B). For (B-C), groups that share a letter are not statistically significantly different (*p* > 0.05) after Tukey’s correction for multiple comparisons. D) Cellular composition of brain lysates for each sample.

Although the ABDK and CD45 MS imparted purer immune cell suspensions than more traditional techniques, a significant CD45-cell population remained (**Figure 1D**). We evaluated two possible sources of this population: non-specific binding due to the concentration of the CD45 magnetic antibody and competition between the CD45 magnetic antibody and the fluorescent CD45 antibody used to identify cells with flow cytometry. To distinguish between these possibilities, we performed a dilution series with the magnetic label and tested several different commercially available fluorescent CD45 antibody clones after MS, collecting enriched and non-enriched fractions to evaluate aberrant retention of different cell populations in different fractions (**Figure S2A**). The gating strategy used to identify broad populations is shown in **Figure S2B**. Cell yield was similar between conditions in the negative fraction and no MS control samples, while yield in the positive fraction seemed to differ based on both experimental conditions (**Figure S2C, Table 2**). Fluorescent epitope labeling should not affect viability due to the speed of the protocol, so the source of this effect is unclear. The microglial proportion in every fraction was not affected by MicroBead dilution, suggesting an equal retention of microglia in the enriched fraction (**Figure S2D, Table 2**). The yield of peripheral myeloid cells and lymphocytes seemed to improve with lower concentrations of MicroBeads, but the aberrant flow-through of these cells into the negative fraction was not affected and the detection of these cells did depend on the fluorescent CD45 antibody clone used (**Figure S2E-F, Table 2**). Crucially, the detected presence of CD45-cells in both positive and negative fractions was influenced only by fluorescent antibody clone used (**Figure S2G, Table 2**). Overall, the purity of samples is demonstrably improved with MS – the detected proportion of immune cells in the enriched fraction is equivalent to or higher than that in non-separated controls (**Figure S2D-G**). We provide some evidence of antibody competition, given that the I3/2-3 CD45 clone seems to improve detection of microglia and lymphocytes while the QA17A26 clone improves detection of non-microglial myeloid cells. In sum, we have demonstrated that the ABDK with CD45 MS reliably captures immune cells from the brain, enriching for cells of interest including lymphocytes and non-microglial myeloid cells.

### CD45 magnetic separation followed by PIPseq captures the complexity of the mouse brain’s immune microenvironment

To demonstrate the utility of our pipeline, we employed a widely used model of neuroinflammation by injecting C57Bl/6J mice with LPS (1.5 x 10^7^ EU/kg, i.p.) or an equivalent volume of sterile saline, and immune cells were isolated from brain tissue with ABDK and CD45 MS 24 hours post-injection. Since the use of enzymes to dissociate brain tissue has been shown to artificially induce immune signatures, mainly in microglia^20,44^, we included inhibitors of transcription, translation, and cell division or an equivalent volume of DMSO vehicle during dissociation as described previously^20^ to examine the extent to which brain immune cells are affected by our dissociation method. After isolation, 40,000 cells per sample were input to capture 20,000 cells in particle-templated instant partitions (PIPs), which were then lysed for downstream cDNA library preparation and sequencing^16^ (**Figure 2A**).

**Figure 2:**
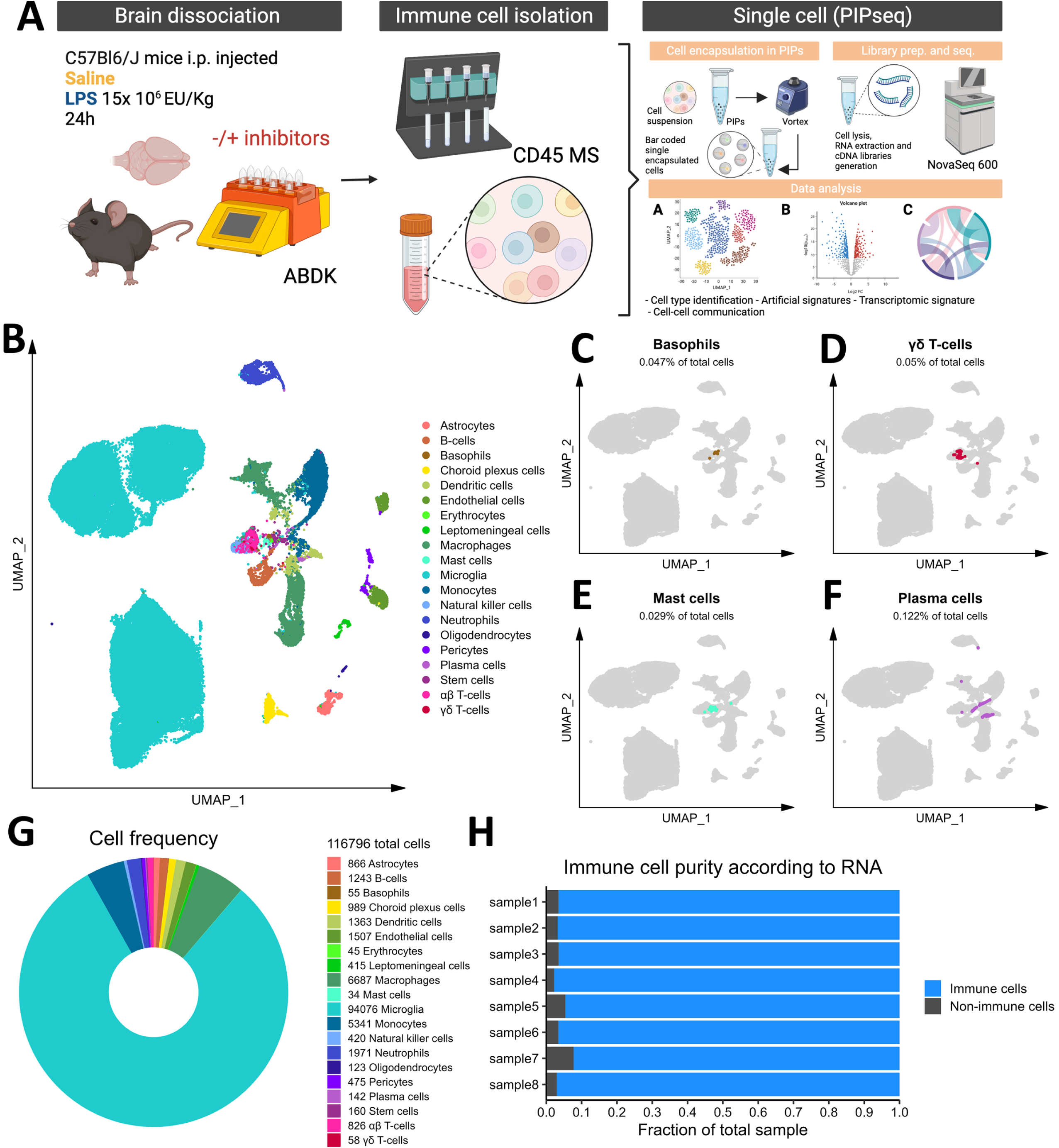
Microfluidics-free single-cell RNA sequencing in brain immune cells. A) Experimental workflow, describing *in vivo* and *ex vivo* treatments and data processing workflow. B) Cellular populations identified, backed by the rationale provided in Figures S3-S9. Basophils (C), mast cells (D), plasma cells (E), and γδ T-cells (F) are highlighted as rare subsets. G) Total cell counts, counts of individual populations, and donut plot showing relative frequencies of our dataset. H) Stacked barchart for each sample showing the relative frequencies of immune cells and non-immune cells.

Initial graph-based clustering revealed 25 populations of cells, which were divided into six broad populations consisting of B-cells, CD45-cells, microglia, myeloid antigen presenting cells (APCs), neutrophils, and T- and natural killer cells (NKs), based on differential expression testing and the expression of canonical markers such as *Ptprc*, *Itgam*, *Ngp*, *Cd3e*, *Ms4a1*, and *Nkg7* (**Figure S3**). We examined each cluster in detail with an iterative subsampling procedure and differential expression testing to confidently annotate cells (**Figure S4** to **Figure S9**). The “B-cell” population was grouped into 6 subtypes (**Figure S4**) that each expressed B-cell markers such as *Cd79a*, *Ms4a1,* and *Iglc3* (**Figure S4D-F**), although one cluster also expressed T-cell genes such as *Cd3e* and *Nkg7* and was designated as a cluster of doublets and removed (**Figure S4G-I**). The CD45-population was grouped into 17 subtypes which expressed markers for astrocytes (e.g. *Aqp4, Gja1, S100b*), choroid plexus cells (e.g. *Folr1*), endothelial cells (e.g. *Cldn5*), leptomeningeal cells (e.g. *Col1a2, Dcn, Slc38a2*), oligodendrocytes (e.g. *Olig1*), and pericytes (e.g. *Acta2*) (Figure S5). The microglial population expressed classic microglial markers such as *Aif1*, *C1qc*, *Hexb*, *P2ry12,* and *Tmem119* (**Figure S6**). The myeloid APCs were clustered into 18 subtypes (**Figure S7A-C**). Several cell types were identified based on marker gene expression, including macrophages (expressing *Mrc1, Cd163, Ms4a7*), monocytes (expressing *Ccr2, Ly6c2, Arg1*), dendritic cells (expressing *Itgax, Cd83, Cd86*), plasma cells (expressing *Igkc, Vpreb3*), and erythrocytes (expressing *Hba-a1*) were identified (**Figure S7D-P**). The neutrophil population consisted of 10 subtypes expressing neutrophil markers such as *Mmp8*, *Ngp*, *S100a8,* or *S100a9* (**Figure S8A-G**). The microglial marker *P2ry12* was expressed in one small cluster, which were classified as doublets and removed (**Figure S8H-I**). Lastly, the T- and NK cell population was grouped into11 subtypes which were annotated as αß T-cells (expressing CD3 alleles, *Cd8b1*), NK cells (expressing *Gzma* and *Klrb1c*), γδ T-cells (expressing γ- and δ-T-cell receptor chains), erythrocytes (expressing *Hba-a1*), macrophages (expressing *Marco*), mast cells (expressing *Mcpt4*), basophils (expressing *Plac8* and *Cebpa*), and stem cells (expressing many immature and proliferating cell markers (**Figure S9**).

In summary, we captured 20 cell types from the mouse brain that consisted of immune cells such as microglia, monocytes, macrophages, B- and T-cells, NKs, dendritic cells, and neutrophils, as well as a some non-immune and/or brain-resident cells (**Figure 2B**). We also identified rare immune populations including basophils, mast cells, plasma cells, and γδ T-cells (**Figure 2C-F**). Our cleaned dataset contained over 116,000 cells from only 8 mouse brains, over 90% of which were immune cell populations (**Figure 2G-H**), demonstrating the efficacy of our FACS-free approach. Together, our data suggest that leveraging automated dissociation, CD45 MS, and the higher cell input granted by PIPseq represents an easily accessible and robust method for identifying immune cells from mouse brain, including rare immune cell populations.

### Myeloid cells are vulnerable to an artificial activation state due to enzymatic dissociation

Enzymes to dissociate the mouse brain have been shown to induce artificial gene activation^20,44^. Hence, we examined to what extent this already defined artificial microglia subpopulation was induced in our pipeline and whether other immune cells were affected. To determine whether the embeddings of cells differed based on inhibitor exposure, we performed a cross-entropy test^29^ on immune cell subsets and the non-immune cell subset. This analysis revealed that granulocytes (composed of the neutrophil, basophil, and mast cell populations), macrophages, microglia, and monocytes were affected by enzymatic dissociation (**Table 3**). The low-dimensional embeddings of each of these populations in the inhibitor-treated and vehicle-treated samples are shown in **Figure 3A-D**. The up-regulation of two published artifactual gene modules^20^ can be seen in cells not treated with inhibitors during dissociation (**Figure 3E-L**). To separate those artifactually activated cells, these three populations were clustered (**Figure 3M-P**), artifact module expression was examined by cluster (**Figure 3Q-T**), and the contribution of each experimental group to each cluster was calculated (**Figure 3U-X**), revealing one cluster of granulocytes, three clusters of macrophages, three clusters of microglia, and one cluster of monocytes that were designated as artifactually activated (**Figure 3Y-BB**). These clusters were removed ahead of biological interrogation of the LPS model.

**Figure 3:**
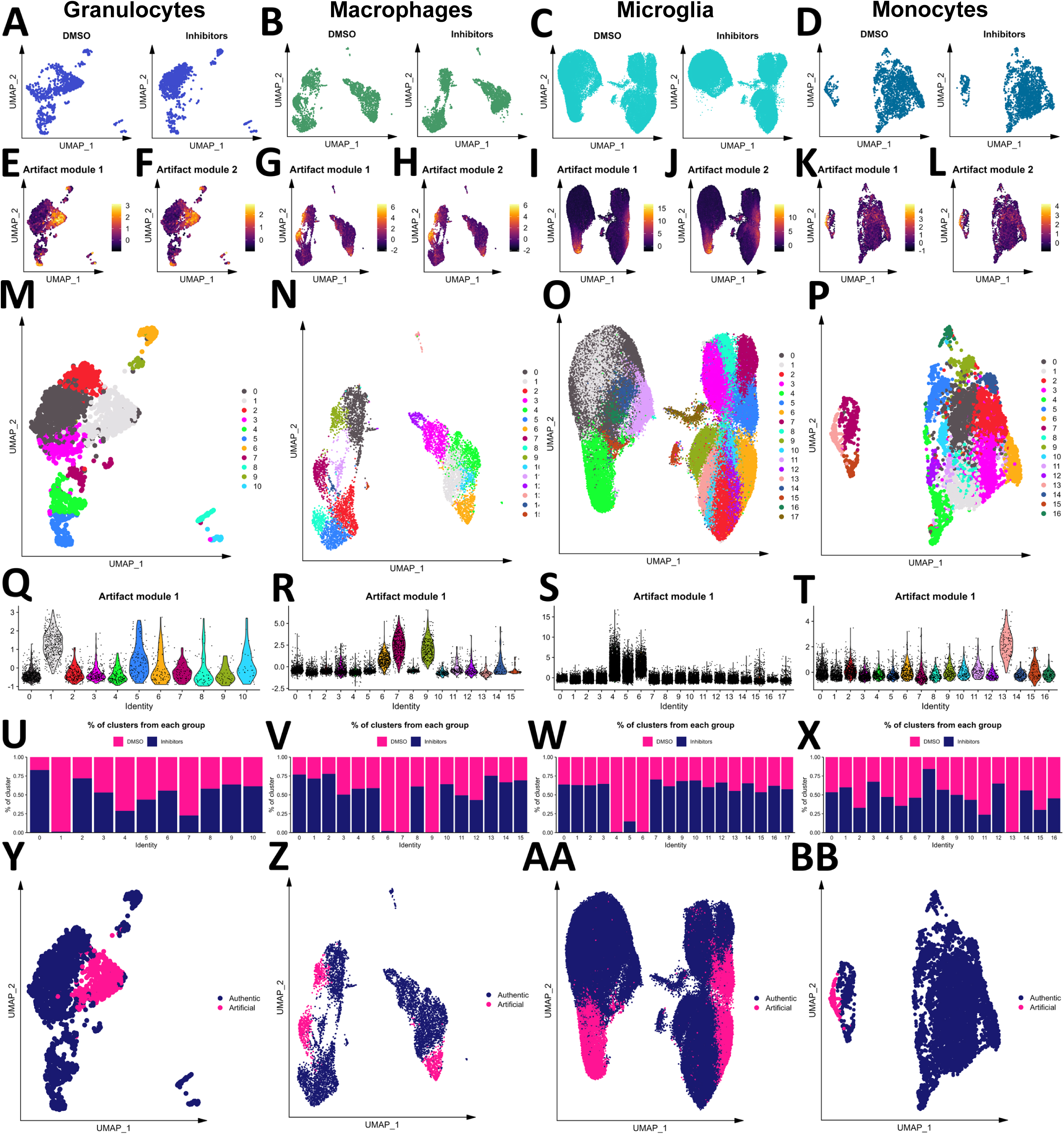
Myeloid cells are vulnerable to enzyme-induced artifactual activation. The embeddings of granulocytes (A), macrophages (B), microglia (C), and monocytes (D) from samples treated with and without inhibitors. Scores of two artifactual gene modules from the literature are shown for granulocytes (E-F), macrophages (G-H), microglia (I-J), and monocytes (K-L). Granulocytes (M), macrophages (N), microglia (O), and monocytes (P) were clustered, and the first artifact module scores were plotted by cluster for each population (Q-T, respectively). The contribution of samples treated with or without inhibitors during dissociation to each granulocyte (U), macrophage (V), microglia (W), and monocyte (X) subset. Final designation of artifact status for granulocytes (Y), macrophages (Z), microglia (AA), and monocytes (BB).

### Peripheral endotoxin activates central-peripheral immune crosstalk

To begin dissecting the unique contributions of immune cells during LPS-induced neuroinflammation, we evaluated whether intraperitoneally administered LPS induces a change in the cellular composition of our brain immune-cell samples. We observed an increase in neutrophils and monocytes and a decrease in B-cells, dendritic cells, and plasma cells in the brain with LPS (**Figure 4A**). The relative abundance of other immune cells and the non-immune cells captured in our pipeline were not significantly affected by LPS (**Figure 4A, Figure S10**). To determine the reason for these changes in immune cell composition with LPS, we inferred the communication of cells using multiple R packages, including CellChat^31^ and MultiNicheNetR^32^. The overall strength of cellular communication was weakened by LPS (**Figure S11A-B**), and the nature of this cellular communication was modulated by LPS (**Figure S11C**) suggesting that circulating endotoxin dysregulates brain immune cell communication. Indeed, the inferred communication between specific pairs of cells is modulated by endotoxin exposure (**Figure 4B-C**). For example, macrophages were expressing fewer ligands with predicted receptors while neutrophils were expressing more ligands (**Figure 4C**). Interestingly, microglia were expressing more ligands whose predicted receptors were expressed by peripheral immune cells, especially monocytes, NKs, and neutrophils, with LPS (**Figure 4C**).

**Figure 4:**
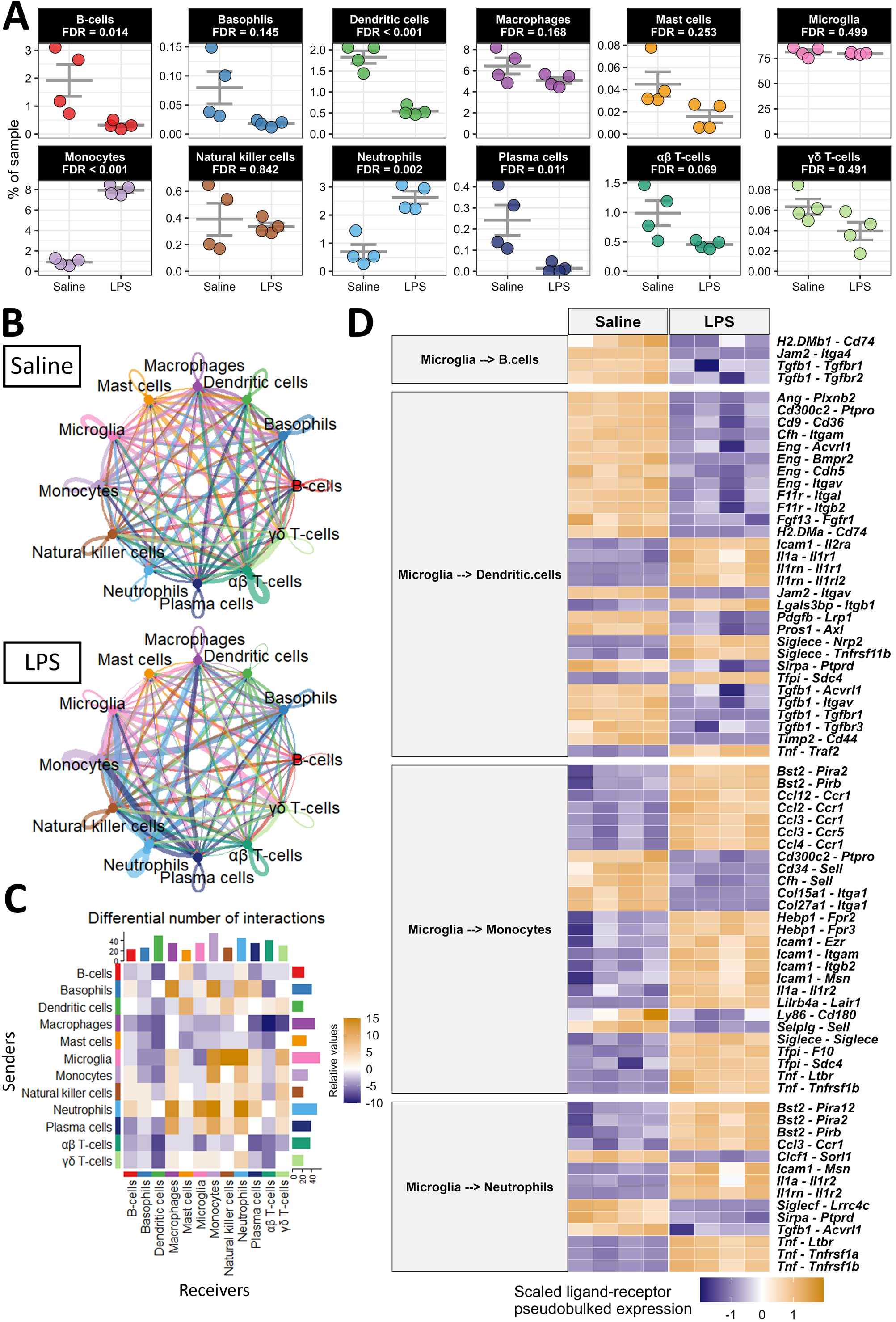
Peripheral LPS exposure activates dynamic central-peripheral immune crosstalk. A) Population frequencies of immune cells from Figure 2 with FDR-adjusted *p*-values from comparing the frequencies in saline- and LPS-treated samples shown for each. B) Circle plot showing the information flow between cells inferred by CellChat for each condition. Lines and bubbles are colored by sender cell type. C) Heatmap showing the differential number of interactions between all combinations of sender (y-axis) and receiver (x-axis) cells. Bars on the top and right show the column-wise and row-wise, respectively, sum of cells. D) Heatmap showing the top 75 ligand-receptor pairs from MultiNicheNet, when microglia were the senders of interest and B-cells, dendritic cells, monocytes, and neutrophils were the receivers of interest. The heatmap is colored by the z-scored pseudobulked expression of the ligand-receptor pairs, and each column is a sample.

We hypothesized that microglia, the brain-resident cells in our dataset, would primarily govern the traffic of cells to and from the brain. We used MultiNicheNetR to predict the specific ligands expressed by microglia and the specific receptors expressed by those immune cells that are differentially frequent in the brain after peripheral LPS exposure. This analysis revealed that chemoattractant and adhesion signals from microglia to B-cells and dendritic cells are largely down-regulated during peripheral LPS exposure. Specifically, several integrins, including *Itgam*, *Itgav*, and *Itgb2* were down-regulated in dendritic cells as were their putative binding partners *Eng*, *F11r*, and *Cfh* in microglia (**Figure 4D**). Communication from microglia to dendritic cells and B-cells through transforming growth factor β (TGFβ) was disrupted by peripheral LPS exposure, and the adhesion of these cells to microglia through major histocompatibility complex class II (MHC-II) receptors (e.g., *H2-DMa, H2-DMb1*, and *Cd74*) and junctional adhesion molecules (JAMs; especially *Jam2*) was also decreased with peripheral LPS exposure (**Figure 4D**). Additionally, we inferred an elevation of several chemoattractant and adhesion signals from microglia to monocytes and neutrophils. The C-C chemokine, interleukin (IL)-1, and tumor necrosis factor (TNF) systems were inferred to be highly active between microglia and these two myeloid populations with peripheral LPS exposure, potentially creating a driving force for the infiltration of these cells. Additionally, the expression of adhesion molecules, especially *Icam1* and *Bst2*, were upregulated in microglia as were their binding partners in monocytes and neutrophils (**Figure 4D**). This analysis revealed novel potential binding partners mediating the extravasation of mononuclear cells in the brain, including the interaction between tetherin (encoded by *Bst2*) and several paired-Ig-like receptors (*Pira2, Pira12,* and *Pirb*), orthologous to human leukocyte immunoglobulin-like receptors, and the interaction between heme binding protein 1 (*Hebp1*) and several formyl peptide receptors (*Fpr2* and *Fpr3*). CellChat analysis corroborates the cell-specific modulation of signaling classes during peripheral LPS exposure. CellChat inferred a loss of incoming JAM and TGFβ and other adhesion molecules in dendritic cells and B-cells during peripheral LPS exposure, while monocytes and neutrophils were predicted to be receiving more interleukins, adhesion ligands, and chemokines (**Figure S11D**). In agreement with the above results, microglia were again predicted to be sending less TGFβ, more C-C chemokines, more TNF, and different adhesion factors (**Figure S11E**).

Finally, we examined the potential loss of communication within microglia inferred by CellChat (**Figure 4C**) using MultiNicheNetR. Of the top 30 differentially regulated ligand-receptor pairs, 26 were down-regulated during LPS, affirming that microglial crosstalk was disrupted with LPS exposure (**Figure S12**). Specifically, peripheral LPS exposure induced a loss of TGFβ signaling, integrin binding, and adhesion molecule expression in microglia. Overall, peripheral LPS exposure disrupted the inferred signaling between microglia, antigen presenting cells, and other microglia. Instead, microglia expressed ligands for receptors highly expressed in peripheral myeloid cells to a greater extent, driving the extravasation of neutrophils and monocytes to the brain during peripheral inflammation.

### LPS induces shared and cell-type specific transcriptomic programs in the brain

As the mobilization and communication of immune cells in the peripheral LPS exposure model is complex, we aimed to distinguish how different cell types contribute to peripheral LPS-induced neuroinflammation. We performed differential expression and gene set enrichment analyses in each cell type by creating pseudo-bulked datasets at the sample level and employing standard bulk RNA sequencing tools. Microglia showed the greatest number of differentially expressed genes (DEGs), while adaptive immune cells and small granulocyte populations showed few DEGs, and peripheral phagocytes regulated a middling number of DEGs (**Figure 5A-B**). We illustrate how many of these DEGs were unique to their respective cell type or shared by at least one other, showing a range of transcriptomic overlap depending on cell type (**Figure 5A-B**).

**Figure 5:**
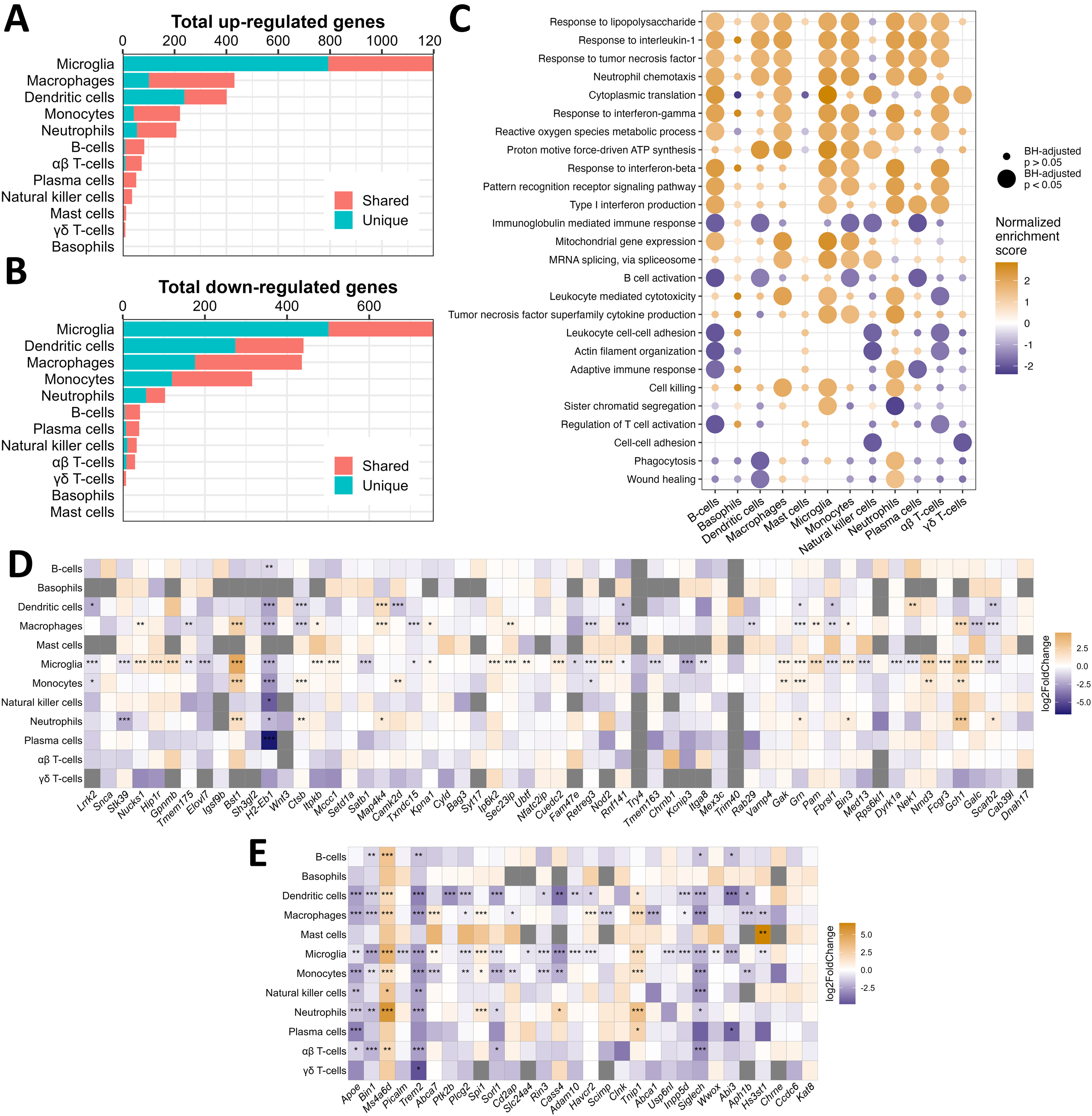
Unique and shared transcriptomic programs in all immune cells dissociated from brain in response to peripheral endotoxin exposure. A) Bar chart showing the total number of genes with adjusted *p* < 0.05 and log_2_ fold change > 2 on pseudobulked datasets. Bar colors show the number of DEGs that are shared by at least one other cell type or unique to that cell type. B) Same as A but showing genes that have log_2_ fold change < −2. C) Dot plot showing the enrichment scores from select GO: BP pathways in all immune cell types after pseudobulked differential expression analysis and gene ranking. Dots are sized according to whether the enrichment met statistical significance (adjusted *p* < 0.05) and colored according to the normalized enrichment score. D) Heatmap showing the log_2_ fold change in expression from saline to LPS of the mouse orthologs of genes associated with the heritability of PD. The x-axis is ranked by ascending *p*-value from ref. ^42^. E) Heatmap showing the log_2_ fold change in expression from saline to LPS of the mouse orthologs of genes associated with the heritability of AD. The x-axis is ranked by ascending *p*-value. This list is borrowed from ref. ^40^ and ref. ^41^. \**p* < 0.05, \*\**p* < 0.01, \*\*\**p* < 0.001

To infer the functional state of each cell type, we employed the well-established gene set enrichment analysis (GSEA)^35,36^ on genes ranked by their differential expression due to peripheral LPS exposure. Several gene sets implicated in LPS signaling, cytokine response, protein synthesis, and metabolic activation were enriched in multiple cell types, suggesting that many immune populations in the brain were metabolically and immunologically responsive to peripheral LPS exposure (**Figure 5C**). However, gene sets associated with cytotoxicity, including the production of TNF, were enriched only in macrophages, monocytes, microglia, and neutrophils, while gene sets associated with cellular adhesion, cytoskeletal organization, and adaptive immune activation were disrupted in B-cells, T-cells, and NKs (**Figure 5C**). Interestingly, neutrophils upregulated genes associated with phagocytosis and wound healing, while microglia and other peripheral phagocytes downregulated or showed no change in these pathways (**Figure 5C**). These data suggest that the neuroimmune profile in the brain induced by peripheral LPS exposure is a product of the activity of many distinct immune cells, and certain inflammatory processes within this profile may be dominated by the activity of peripheral immune cells.

Finally, we curated lists of risk factors identified for Parkinson’s disease (PD) and Alzheimer’s disease (AD) in meta-genome wide association studies (meta-GWAS) and examined the change in expression of these genes in each cell type due to an inflammatory stimulus. Microglia regulated many of these genes in response to peripheral LPS exposure, and some genes are uniquely regulated by microglia, including *Gpnmb*, *Tmem163*, *Picalm*, and *Slc2a4* (**Figure 5D-E**). However, many of these genes were also regulated by peripheral myeloid cells in response to peripheral LPS exposure, including *Bst1*, *Gch1*, *Lrrk2*, *Satb1*, *Apoe*, *Plcg2*, *Siglech*, *Sorl1*, *Spi1* (**Figure 5D-E**). Most immune populations captured in this study up-regulated *Ms4a6d* and *Tnip1* and downregulated *H2-Eb1*, *Bin1,* and *Trem2* after peripheral LPS exposure (**Figure 5D-E**). Interestingly, certain genes differed in the direction of their regulation between microglia and peripheral immune cells in response to peripheral LPS exposure, including *Galc*, *Nek1*, *Abca7*, and *Cass4*, while only peripheral populations regulated genes such as *Camk2d*, *Ctsb*, *Grn*, *Map4k4*, *Abca1*, *Aph1b*, *Cd2ap*, and *Ptk2b*. Overall, our findings reveal the existence of immune cell subset-specific expression of genes associated with risk for neurodegeneration in response to peripheral LPS, consistent with the growing perspective that genes that confer risk to neurodegenerative disease are linked to the immune system.

## Discussion

In this study, we report a readily accessible, robust, and reliable experimental approach to interrogate the mouse brain’s immune system. First, we compared the efficacy of competing protocols designed to isolate immune cells from the mouse brain and demonstrated that automated dissociation and MS yields purer samples and greater cell numbers compared to Percoll separation after collagenase digestion. Second, we demonstrated the efficacy of the recent microfluidics-free scRNAseq platform PIPseq^16^ combined with automated dissociation and CD45 MS in capturing the complexity of the brain’s immune system, including changes induced by peripheral LPS exposure.

It is important to note that the estimated fraction of immune cells from this strategy differed between flow cytometry and scRNAseq analyses, where the purity of immune cells was much greater according to RNA than surface epitopes. The cleavage of surface epitopes by enzymes during dissociation is a documented disadvantage, which likely interferes with the labeling of key cell markers including CD45 and CD11b^44^. Additionally, other reports using Percoll-mediated separation reported more successful immune enrichment as measured by flow cytometry than what we obtained here^43^, but differences in the enzyme cocktails used may partially underlie this discrepancy. Indeed, the immune purity suggested by RNA is similar between our studies. In sum, if surface labeling is a desired downstream application, enzyme-free methodologies coupled with MS may be more advantageous.

We replicated previous findings that enzymatic dissociation can induce false gene signatures in microglia^20,44^, which must be considered with care when interrogating the brain’s immune response to stimuli. The artifactual gene signature, comprised mostly of immediate early genes, heat shock protein genes, and some chemokines and cytokines^20^, was well preserved in our study. However, in addition to microglia, we note the induction of this false signature in other myeloid cells, including neutrophils, macrophages, and monocytes, making the proper control over this false signature even more pressing. Importantly, the pharmacological inhibition of this genomic artifact established previously^20^ was also well preserved in our study, enabling an artifact-free query into brain-immune interactions.

In the peripheral LPS model, we observed the trafficking of neutrophils to the brain, which is a documented effect of peripheral endotoxin administration. The influx of neutrophils due to peripheral LPS exposure has been confirmed by others using immunohistochemical approaches^45^ and flow cytometry techniques^18,46^. *In vivo* two-photon imaging has revealed physical contact between microglia and neutrophils in the brain of peripheral LPS-treated mice^47^, consistent with the heightened communication between microglia and neutrophils we report here, especially the increased expression of adhesion molecules, such as *Icam1* and *Msn* respectively, on these two cell types. We also demonstrated the increased expression of IL-1 receptors, TNF receptors, and C-C chemokine receptors on neutrophils, which have been shown to induce the migration of neutrophils^48–50^ and are likely enabling a downstream contact with microglia.

The trafficking of monocytes to the brain due to peripheral LPS, another established consequence of the model and other models of peripheral inflammation, was also observed here. Fate mapping with reporter mouse models^51^ and flow cytometry^19^ have demonstrated the migration of monocytes to the brain during LPS exposure the necessity of CCL2 in inducing this traffic. Our single-cell data support the induction of *Ccl2* signaling from microglia to monocytes after peripheral LPS exposure, but our analysis suggests that several other microglial C-C chemokine ligands, including CCL3 and CCL4, and monocyte C-C chemokine receptors, including CCR1 and CCR5, may also be active. Monocytic infiltration of the brain has been reported in a model of hepatic inflammation, where TNF was implicated as a key regulator of this traffic^52^. Our intercellular communication analyses support the role of TNF, and in recent work we observed increased monocytes in the brain of animals that experienced colonic inflammation induced by dextran sulfate sodium^53^. Moreover, monocytes have been implicated as key disease effectors in models of central nervous system damage including models of ischemia, neurodegeneration, and demyelination^54^, suggesting that the relationship we report here between brain resident and peripheral phagocytes has broad disease implications. Additionally, physical contact between microglia and infiltrating monocytes triggered by peripheral inflammation has not been reported, but the induction of adhesion systems between these cell types we report warrants the inspection of this relationship and its consequences on brain health.

In addition, our scRNAseq approach reveals the unique and overlapping contributions of different cell types to the broad effects of peripheral inflammation that have been reported previously. Peripheral LPS exposure induces interferon signaling in the brain which is generally attributed to microglia^55^, but our single-cell approach suggests that interferon production and signaling is induced in both innate and adaptive immune cells in the brain, in line with other reports from non-CNS and *ex vivo* systems^56–59^. TGFβ signaling between brain cells is increased during aging and ischemia^60^ which we posit is a neuroprotective mechanism^61^. It is known that LPS antagonizes TGFβ in microglia^62^, but we report that microglial TGFβ signaling to peripheral immune cells is also disrupted after peripheral LPS exposure, suggesting that peripheral inflammation dysregulates homeostatic immune crosstalk. Leukocyte adhesion, which we found is increased between microglia and neutrophils via *Bst2* and *Icam1* and microglia and monocytes via *Icam1, Bst2*, and several integrins, but dysregulated in NKs and adaptive immune cells, has not been directly examined in the brain under conditions of peripheral inflammation, but it has been observed in the human lymphatic system^63^, *in vitro* tumor systems^64,65^, and mouse kidney^66^ after LPS exposure. Finally, we suggest that several non-microglial immune cell types contribute to the classic readouts of peripheral LPS-induced neuroinflammation, including TNF production, interleukin production, and metabolic activation^67^, a finding that has implications for future cell type-specific targeting in therapeutic interventions.

Further, the mobilization of disease-related genes in the brain in response to peripheral inflammation shows complex cell-type specificity. Specifically, we found that genes implicated in the heritability of PD and AD were regulated by microglia during peripheral inflammation, supporting the growing evidence that the pathogenesis of these diseases is linked to immune function^68–71^. However, many of these genes were regulated by peripheral immune cells, either in addition to or differently from microglia, suggesting that more intense focus needs to be placed on cells outside the brain. This perspective is beginning to be adopted by the field with respect to genes implicated in the endolysosomal system including leucine-rich repeat kinase (LRRK2)^72^, progranulin (GRN), and glycoprotein non-metastatic B (GPNMB) as risk genes for several neurodegenerative diseases^3,73,74^. Interestingly, disease-related genes were regulated largely by myeloid cells in our study. Yet it is known that genomic programs characterized by the same genes may be activated in T-cells during aging^41^ and vascular cells during AD^40^, suggesting that the cell-type specificity of risk-gene expression we report here may be more complex in the chronically inflamed brain or disease states.

Due to the CD45 MS used to enhance the immune cell purity of our samples, we captured only small amounts of astrocytes, pericytes, and endothelial cells in our scRNAseq dataset, which limits the insight we can gain about the role of these cells during peripheral LPS-induced neuroinflammation. The neurovascular unit, comprised of these cells, is a key intermediary in central-peripheral immune crosstalk. LPS may be injurious to the blood brain barrier (BBB) in certain doses and conditions^75^, and leukocytes adhere to the endothelium before entering the brain parenchyma^76^. In our study, we cannot discern whether monocytes and neutrophils are infiltrating due to increased cellular adhesion with endothelial cells or to BBB damage. We also cannot determine if microglial chemoattractant signals are reaching these peripheral myeloid cells directly or astrocytes and endothelial cells, which are known producers of chemoattractants and interleukins^77,78^, are mediating this crosstalk. Yet, unraveling the complexity of this multicellular system is tractable proposition by using FACS or MS procedures to isolate both immune and neurovascular cells coupled with the scalability offered by PIPseq^16^.

In summary, we have demonstrated the utility of an accessible and robust platform to delineate the unique contributions of brain-resident and extravasating immune cells to peripheral endotoxin-induced neuroinflammation. Our single-cell genomics approach suggests that microglia preferentially communicate with peripheral myeloid cells rather than other microglia after peripheral LPS exposure, which may have far-reaching consequences for maintaining brain health^79^. The nature of this central-peripheral immune crosstalk in other animal models of neuroinflammation represents a source of untapped insight for human disease that will be feasible to interrogate using our methodology. Whether microglia are mobilizing the adaptive immune system under conditions of chronic peripheral inflammation directly or indirectly through long-term communication with peripheral antigen-presenting cells is a therapeutically relevant area^80,81^. Finally, whether innate immune cells are recruited to the brain in a similar manner in other cases of acute inflammation is also of interest to the field, as is the extent to which abrogation of this recruitment is beneficial for brain health. These lines of study, enabled by the experimental approach described herein, may unlock new targets to improve brain health through modulation of central-peripheral neuroimmune crosstalk.

## AUTHORS CONTRIBUTIONS

JSB, OUH and MGT designed the study. JSB and OUH performed the experiments. JSB performed statistical and bioinformatic analysis and created figures. JSB, OUH and MGT analyzed the results and wrote the manuscript. All authors read and approved the final manuscript.

## Acknowledgements

We thank Rebecca Wallings and Maeve Krueger for assistance in editing the manuscript, and the whole of the Tansey lab for useful discussions. Funding for this work was derived from a Parkinson’s Foundation Research Center of Excellence award (MGT), the Michael J. Fox Foundation for Parkinson’s Research (MGT), and the National Institutes of Health (RF1NS128800, MGT) and the Interdisciplinary Training in Movement Disorders and Neurorestoration T32 at the University of Florida (T32-NS082128, JSB).

## Tables

Table 1: Results from two-way ANOVAs demonstrating the main effect of genotype, main effect of isolation protocol, and interaction on the raw counts and relative frequencies of each immune cell population. These statistics accompany the experiment presented in Figure 1.

Table 2: Results from ANOVAs demonstrating the effects of MicroBead dilution and CD45 antibody clone on immune cell detection after brain dissociation. These statistics accompany the experiment presented in Figure S2.

Table 3: Results from a cross-entropy test in every cell subset highlighted in Figure 2.

Table S1: Product information for antibodies used to compare the efficacy of different brain immune cell isolation techniques.

Table S2: Product information for antibodies used to evaluate whether magnetic antibody labeling interferes with downstream flow cytometry.

## Figure Captions

Figure S1: Gating strategy and raw count data for Figure 1. A) Gating strategy used to identify cell subsets. Raw counts of T-cells (B), B-cells (C), CD45-cells (D), dendritic cells (E), neutrophils (F), microglia (G), and monocytes/macrophages (H) are shown. For each plot, groups that share a letter are not statistically significantly different (*p* > 0.05) after Tukey’s correction for multiple comparisons.

Figure S2: Testing different MicroBead dilutions and CD45 antibody clones in the detected yields of immune cells from brain. A) Experimental workflow showing the sample fractions collected after magnetic separation and CD45 antibody clones used during surface labeling. B) Gating strategy used to identify broad cell subsets. C) Live cell count in each fraction. The frequency of microglia (D), non-microglia myeloid cells (E), lymphocytes (F), and CD45-cells (G) are shown, represented as a percentage of total live cells in the sample. In these figures, groups that share a letter are not statistically significant (*p* > 0.05) after Tukey’s correction for multiple comparisons, which were conducted separately for each fraction.

Figure S3: Initial clustering and annotation of our brain immune PIPseq dataset. A) Cells were clustered with a low resolution at first, leaving 25 clusters, labeled on a UMAP plot. The expression of *Ptprc* (B), *Itgam* (C), *Ngp* (D), *Cd3e* (E), *Ms4a1* (F), *Nkg7* (G) is shown, expressed as log2-transformed normalized counts. H) Clusters were labeled broadly based on the expression of canonical cell markers. These clusters were then isolated and analyzed in more depth in later figures, shown in (I-N).

Figure S4: Annotation of the B-cell cluster. A) Robust clustering revealed 6 groups of cells. The effectiveness of this clustering strategy is shown by the silhouette score in each cell (B) and average co-clustering of cells during iterative subsampling (C). Useful genes in cluster identification were *Cd79a* (D), *Ms4a1* (E), *Ighm* (F), *Cd3e* (G), and *Nkg7* (H). I) Final cluster identities included B-cells and a doublet cluster, which showed expression of T- and B-cell markers, that was removed before further analyses.

Figure S5: Annotation of the CD45-cluster. A) Robust clustering revealed 17 groups of cells. The effectiveness of this clustering strategy is shown by the silhouette score of each cell (B) and average co-clustering of cells during iterative subsampling (C). Useful genes in cluster identification were *Acta2* (D), *Aqp4* (E), *Cldn5* (F), *Col1a2* (G), *Dcn* (H), *Folr1* (I), *Gja1* (J), *Olig1* (K), *S100b* (L), and *Slc38a2* (M). N) Final cluster identities included choroid plexus cells, endothelial cells, astrocytes, pericytes, leptomeningeal cells, and oligodendrocytes.

Figure S6: Annotation of the microglial cluster. A) Robust clustering revealed 7 clusters in a down-sampled version of our microglial dataset. The effectiveness of this clustering strategy is shown by the silhouette score in each cell (B) and average co-clustering of cells during iterative subsampling (C). D) The clustering resolution indicated by chooseR was applied to the full dataset, leaving 14 clusters. The Dice coefficient was calculated for each pair of down-sampled and unsampled clusters, plotted as a heatmap. Unsampled clusters are shown in (E). Microglial marker genes are shown, including *Aif1* (F), *C1qc* (G), *Hexb* (H), *P2ry12* (I), *Tmem119* (J).

Figure S7: Annotation of the myeloid antigen-presenting cell cluster. A) Robust clustering revealed 16 clusters. The effectiveness of this clustering strategy is shown by the silhouette score of each cell (B) and average co-clustering of cells during iterative subsampling (C). Useful marker genes in cluster annotation were *Mrc1* (D), *Cd163* (E), *Ms4a7* (F), *Itgax* (G), *Cd83* (H), *Cd86* (I), *Ccr2* (J), *Ly6c2* (K), *Arg1* (L), *Igkc* (M), *Vpreb3* (N), and *Hba-a1* (O). P) Final cluster identities were monocytes, macrophages, dendritic cells, plasma cells, and erythrocytes.

Figure S8: Annotation of the neutrophil cluster. A) Robust clustering revealed 10 clusters. The effectiveness of this clustering strategy is shown by the silhouette score of each cell (B) and average co-clustering of cells during iterative subsampling (C). Useful marker genes in cluster identification were *Mmp8* (D), *Ngp* (E), *S100a8* (F), *S100a9* (G), and *P2ry12* (H). In total, this dataset contained neutrophils and a small cluster of doublets, marked by the expression of microgial and neutrophil genes, which were removed prior to further analyses.

Figure S9: Annotation of the T- and natural killer cell cluster. A) Robust clustering revealed 12 clusters. The effectiveness of this clustering strategy is shown by the silhouette score of each cell (B) and average co-clustering of cells during iterative subsampling (C). Useful marker genes in cluster annotation were *Cd3e* (D), *Cd8b1* (E), *Gzma* (F), *Klrb1c* (G), *Trdc* (H), *Trgc1* (I), *Marco* (J), *Hba-a1* (K), *Cebpa* (L), *Mcpt4* (M), *Plac8* (N), and *Calca* (O). P) Final identities were αβ T-cells, natural killer cells, stem cells, γδ T-cells, basophils, macrophages, mast cells, and erythrocytes.

Figure S10: Population frequencies of non-immune cells from Figure 2, with FDR-adjusted *p*-values shown for each.

Figure S11: Intercellular immune signaling is affected by peripheral endotoxin. A) Bar chart showing the total number of cellular interactions inferred by CellChat. B) Bar chart showing the relative weight of all cellular interactions by treatment group inferred by CellChat. C) Bar chart showing the number of interactions within each signaling class included in the CellChatDB database, scaled to range from 0 to 1. D) Heatmap showing the relative strength of incoming signals (i.e., receptor expression) within each CellChat signaling class, separated by cell type on the x-axis within each treatment. Bars on the top and right show the aggregated relative strength of receivers and signaling classes, respectively. E) Heatmap showing the relative strength of outgoing signals (i.e., ligand expression) within each CellChat signaling class, separated by cell type on the x-axis within each treatment. Bars on the top and right show the aggregated relative strength of senders and signaling class, respectively.

Figure S12: Peripheral LPS exposure disrupts microglial communication. The top 30 ligand-receptor pairs between microglia inferred from MultiNicheNet are shown on the right y-axis. The heatmap is colored by the z-scored pseudobulked expression of the ligand-receptor pairs, and each column is a sample.

## References

1. Akiyama, H. et al. Inflammation and Alzheimer’s disease. Neurobiol. Aging 21, 383–421 (2000).

2. Berk, M. et al. So depression is an inflammatory disease, but where does the inflammation come from? BMC Med. 11, 200 (2013).

3. Tansey, M. G. et al. Inflammation and immune dysfunction in Parkinson disease. Nat. Rev. Immunol. 22, 657–673 (2022).

4. Hammond, T. R., Robinton, D. & Stevens, B. Microglia and the brain: complementary partners in development and disease. Annu. Rev. Cell Dev. Biol. 34, 523–544 (2018).

5. Kannarkat, G. T., Boss, J. M. & Tansey, M. G. The role of innate and adaptive immunity in Parkinson’s disease. J Parkinsons Dis 3, 493–514 (2013).

6. Hammond, T. R. et al. Single-Cell RNA Sequencing of Microglia throughout the Mouse Lifespan and in the Injured Brain Reveals Complex Cell-State Changes. Immunity 50, 253–271.e6 (2019).

7. Masuda, T., Sankowski, R., Staszewski, O. & Prinz, M. Microglia Heterogeneity in the Single-Cell Era. Cell Rep. 30, 1271–1281 (2020).

8. Perfetto, S. P., Chattopadhyay, P. K. & Roederer, M. Seventeen-colour flow cytometry: unravelling the immune system. Nat. Rev. Immunol. 4, 648–655 (2004).

9. Delaney, C. et al. Combinatorial prediction of marker panels from single-cell transcriptomic data. Mol. Syst. Biol. 15, e9005 (2019).

10. Rundberg Nilsson, A., Bryder, D. & Pronk, C. J. H. Frequency determination of rare populations by flow cytometry: a hematopoietic stem cell perspective. Cytometry A 83, 721–727 (2013).

11. Spitzer, M. H. & Nolan, G. P. Mass cytometry: single cells, many features. Cell 165, 780–791 (2016).

12. Cossarizza, A. et al. Guidelines for the use of flow cytometry and cell sorting in immunological studies. Eur. J. Immunol. 47, 1584–1797 (2017).

13. Efremova, M., Vento-Tormo, R., Park, J.-E., Teichmann, S. A. & James, K. R. Immunology in the Era of Single-Cell Technologies. Annu. Rev. Immunol. 38, 727–757 (2020).

14. Li, X. et al. Single-cell transcriptomic analysis of the immune cell landscape in the aged mouse brain after ischemic stroke. J. Neuroinflammation 19, 83 (2022).

15. Chen, X. et al. Microglia-mediated T cell infiltration drives neurodegeneration in tauopathy. Nature 615, 668–677 (2023).

16. Clark, I. C. et al. Microfluidics-free single-cell genomics with templated emulsification. Nat. Biotechnol. (2023) doi:10.1038/s41587-023-01685-z.

17. Hatori, M. N., Kim, S. C. & Abate, A. R. Particle-Templated Emulsification for Microfluidics-Free Digital Biology. Anal. Chem. 90, 9813–9820 (2018).

18. Aguilar-Valles, A., Kim, J., Jung, S., Woodside, B. & Luheshi, G. N. Role of brain transmigrating neutrophils in depression-like behavior during systemic infection. Mol. Psychiatry 19, 599–606 (2014).

19. Cazareth, J., Guyon, A., Heurteaux, C., Chabry, J. & Petit-Paitel, A. Molecular and cellular neuroinflammatory status of mouse brain after systemic lipopolysaccharide challenge: importance of CCR2/CCL2 signaling. J. Neuroinflammation 11, 132 (2014).

20. Marsh, S. E. et al. Dissection of artifactual and confounding glial signatures by single-cell sequencing of mouse and human brain. Nat. Neurosci. 25, 306–316 (2022).

21. Dobin, A. et al. STAR: ultrafast universal RNA-seq aligner. Bioinformatics 29, 15–21 (2013).

22. Hao, Y. et al. Integrated analysis of multimodal single-cell data. Cell 184, 3573–3587. (2021).

23. Stuart, T. et al. Comprehensive Integration of Single-Cell Data. Cell 177, 1888–1902.e21 (2019).

24. Ahlmann-Eltze, C. & Huber, W. glmGamPoi: fitting Gamma-Poisson generalized linear models on single cell count data. Bioinformatics 36, 5701–5702 (2021).

25. Hu, C. et al. CellMarker 2.0: an updated database of manually curated cell markers in human/mouse and web tools based on scRNA-seq data. Nucleic Acids Res. 51, D870–D876 (2023).

26. Patterson-Cross, R. B., Levine, A. J. & Menon, V. Selecting single cell clustering parameter values using subsampling-based robustness metrics. BMC Bioinformatics 22, 39 (2021).

27. Phipson, B. et al. propeller: testing for differences in cell type proportions in single cell data. Bioinformatics 38, 4720–4726 (2022).

28. Ritchie, M. E. et al. limma powers differential expression analyses for RNA- sequencing and microarray studies. Nucleic Acids Res. 43, e47 (2015).

29. Roca, C. P. et al. A cross entropy test allows quantitative statistical comparison of t-SNE and UMAP representations. Cell Rep. Methods 3, 100390 (2023).

30. Bunis, D. G., Andrews, J., Fragiadakis, G. K., Burt, T. D. & Sirota, M. dittoSeq: universal user-friendly single-cell and bulk RNA sequencing visualization toolkit. Bioinformatics 36, 5535–5536 (2021).

31. Jin, S. et al. Inference and analysis of cell-cell communication using CellChat. Nat. Commun. 12, 10881 (2021).

32. Browaeys, R., et al. MultiNicheNet: a flexible framework for differential cell-cell communication analysis from multi-sample multi-condition single-cell transcriptomics data. BioRxiv (2023) doi:10.1101/2023.06.13.544751.

33. Crowell, H. L. et al. muscat detects subpopulation-specific state transitions from multi-sample multi-condition single-cell transcriptomics data. Nat. Commun. 11, 6077 (2020).

34. Love, M. I., Huber, W. & Anders, S. Moderated estimation of fold change and dispersion for RNA-seq data with DESeq2. Genome Biol. 15, 550 (2014).

35. Subramanian, A. et al. Gene set enrichment analysis: a knowledge-based approach for interpreting genome-wide expression profiles. Proc Natl Acad Sci USA 102, 15545–15550 (2005).

36. Wu, T. et al. clusterProfiler 4.0: A universal enrichment tool for interpreting omics data. Innovation (Camb) 2, 100141 (2021).

37. Ashburner, M. et al. Gene Ontology: tool for the unification of biology. Nat. Genet. 25, 25–29 (2000).

38. Gene Ontology Consortium et al. The Gene Ontology knowledgebase in 2023. Genetics 224, (2023).

39. Zhu, A., Ibrahim, J. G. & Love, M. I. Heavy-tailed prior distributions for sequence count data: removing the noise and preserving large differences. Bioinformatics 35, 2084–2092 (2019).

40. Yang, A. C. et al. A human brain vascular atlas reveals diverse mediators of Alzheimer’s risk. Nature 603, 885–892 (2022).

41. Piehl, N. et al. Cerebrospinal fluid immune dysregulation during healthy brain aging and cognitive impairment. Cell 185, 5028–5039.e13 (2022).

42. Nalls, M. A. et al. Identification of novel risk loci, causal insights, and heritable risk for Parkinson’s disease: a meta-analysis of genome-wide association studies. Lancet Neurol. 18, 1091–1102 (2019).

43. Guldner, I. H., Golomb, S. M., Wang, Q., Wang, E. & Zhang, S. Isolation of mouse brain-infiltrating leukocytes for single cell profiling of epitopes and transcriptomes. STAR Protocols 2, 100537 (2021).

44. Mattei, D. et al. Enzymatic dissociation induces transcriptional and proteotype bias in brain cell populations. Int. J. Mol. Sci. 21, (2020).

45. Jeong, H.-K., Jou, I. & Joe, E. Systemic LPS administration induces brain inflammation but not dopaminergic neuronal death in the substantia nigra. Exp. Mol. Med. 42, 823–832 (2010).

46. Byun, D. J., Lee, J., Yu, J.-W. & Hyun, Y.-M. NLRP3 Exacerbate NETosis-Associated Neuroinflammation in an LPS-Induced Inflamed Brain. Immune Netw. 23, e27 (2023).

47. Kim, Y. R. et al. Neutrophils Return to Bloodstream Through the Brain Blood Vessel After Crosstalk With Microglia During LPS-Induced Neuroinflammation. Front. Cell Dev. Biol. 8, 613733 (2020).

48. Vieira, S. M. et al. A crucial role for TNF-alpha in mediating neutrophil influx induced by endogenously generated or exogenous chemokines, KC/CXCL1 and LIX/CXCL5. Br. J. Pharmacol. 158, 779–789 (2009).

49. Montecucco, F. et al. Tumor necrosis factor-alpha (TNF-alpha) induces integrin CD11b/CD18 (Mac-1) up-regulation and migration to the CC chemokine CCL3 (MIP-1alpha) on human neutrophils through defined signalling pathways. Cell. Signal. 20, 557–568 (2008).

50. Pyrillou, K., Burzynski, L. C. & Clarke, M. C. H. Alternative Pathways of IL-1 Activation, and Its Role in Health and Disease. Front. Immunol. 11, 613170 (2020).

51. Chen, H.-R. et al. Monocytes promote acute neuroinflammation and become pathological microglia in neonatal hypoxic-ischemic brain injury. Theranostics 12, 512–529 (2022).

52. D’Mello, C., Le, T. & Swain, M. G. Cerebral microglia recruit monocytes into the brain in response to tumor necrosis factoralpha signaling during peripheral organ inflammation. J. Neurosci. 29, 2089–2102 (2009).

53. Boles, J. S., et al. A leaky gut dysregulates gene networks in the brain associated with immune activation, oxidative stress, and myelination in a mouse model of colitis. BioRxiv (2023) doi:10.1101/2023.08.10.552488.

54. Spiteri, A. G., Wishart, C. L., Pamphlett, R., Locatelli, G. & King, N. J. C. Microglia and monocytes in inflammatory CNS disease: integrating phenotype and function. Acta Neuropathol. 143, 179–224 (2022).

55. Lively, S. & Schlichter, L. C. Microglia Responses to Pro-inflammatory Stimuli (LPS, IFNγ+TNFα) and Reprogramming by Resolving Cytokines (IL-4, IL-10). Front. Cell. Neurosci. 12, 215 (2018).

56. Jacobs, A. T. & Ignarro, L. J. Lipopolysaccharide-induced expression of interferon-beta mediates the timing of inducible nitric-oxide synthase induction in RAW 264.7 macrophages. J. Biol. Chem. 276, 47950–47957 (2001).

57. Fultz, M. J., Barber, S. A., Dieffenbach, C. W. & Vogel, S. N. Induction of IFN-gamma in macrophages by lipopolysaccharide. Int. Immunol. 5, 1383–1392 (1993).

58. Varma, T. K., Lin, C. Y., Toliver-Kinsky, T. E. & Sherwood, E. R. Endotoxin-induced gamma interferon production: contributing cell types and key regulatory factors. Clin. Diagn. Lab. Immunol. 9, 530–543 (2002).

59. Ali, S. et al. Sources of type I interferons in infectious immunity: plasmacytoid dendritic cells not always in the driver’s seat. Front. Immunol. 10, 778 (2019).

60. Doyle, K. P., Cekanaviciute, E., Mamer, L. E. & Buckwalter, M. S. TGFβ signaling in the brain increases with aging and signals to astrocytes and innate immune cells in the weeks after stroke. J. Neuroinflammation 7, 62 (2010).

61. Kashima, R. & Hata, A. The role of TGF-β superfamily signaling in neurological disorders. Acta Biochim Biophys Sin (Shanghai) 50, 106–120 (2018).

62. Mitchell, K. et al. LPS antagonism of TGF-β signaling results in prolonged survival and activation of rat primary microglia. J. Neurochem. 129, 155–168 (2014).

63. Sawa, Y. et al. LPS-induced IL-6, IL-8, VCAM-1, and ICAM-1 expression in human lymphatic endothelium. J. Histochem. Cytochem. 56, 97–109 (2008).

64. Park, G.-S. & Kim, J.-H. LPS Up-Regulates ICAM-1 Expression in Breast Cancer Cells by Stimulating a MyD88-BLT2-ERK-Linked Cascade, Which Promotes Adhesion to Monocytes. Mol. Cells 38, 821–828 (2015).

65. Bui, T. M., Wiesolek, H. L. & Sumagin, R. ICAM-1: A master regulator of cellular responses in inflammation, injury resolution, and tumorigenesis. J. Leukoc. Biol. 108, 787–799 (2020).

66. Kajiwara, K., Sawa, Y., Fujita, T. & Tamaoki, S. Immunohistochemical study for the expression of leukocyte adhesion molecules, and FGF23 and ACE2 in P. gingivalis LPS-induced diabetic nephropathy. BMC Nephrol. 22, 3 (2021).

67. Batista, C. R. A., Gomes, G. F., Candelario-Jalil, E., Fiebich, B. L. & de Oliveira, A. C. P. Lipopolysaccharide-Induced Neuroinflammation as a Bridge to Understand Neurodegeneration. Int. J. Mol. Sci. 20, (2019).

68. Pierce, S. & Coetzee, G. A. Parkinson’s disease-associated genetic variation is linked to quantitative expression of inflammatory genes. PLoS ONE 12, e0175882 (2017).

69. Sims, R. et al. Rare coding variants in PLCG2, ABI3, and TREM2 implicate microglial-mediated innate immunity in Alzheimer’s disease. Nat. Genet. 49, 1373–1384 (2017).

70. Kannarkat, G. T. et al. Common Genetic Variant Association with Altered HLA Expression, Synergy with Pyrethroid Exposure, and Risk for Parkinson’s Disease: An Observational and Case-Control Study. npj Parkinsons Disease 1, 15002 (2015).

71. Garretti, F. et al. Interaction of an α-synuclein epitope with HLA-DRB1∗15:01 triggers enteric features in mice reminiscent of prodromal Parkinson’s disease. Neuron (2023) doi:10.1016/j.neuron.2023.07.015.

72. Wallings, R. L., Herrick, M. K. & Tansey, M. G. LRRK2 at the interface between peripheral and central immune function in parkinson’s. Front. Neurosci. 14, 443 (2020).

73. Houser, M. C. et al. Progranulin loss results in sex-dependent dysregulation of the peripheral and central immune system. Front. Immunol. 13, 1056417 (2022).

74. Wilson, D. M. et al. Hallmarks of neurodegenerative diseases. Cell 186, 693–714 (2023).

75. Banks, W. A. et al. Lipopolysaccharide-induced blood-brain barrier disruption: roles of cyclooxygenase, oxidative stress, neuroinflammation, and elements of the neurovascular unit. J. Neuroinflammation 12, 223 (2015).

76. Peng, X., Luo, Z., He, S., Zhang, L. & Li, Y. Blood-Brain Barrier Disruption by Lipopolysaccharide and Sepsis-Associated Encephalopathy. Front. Cell. Infect. Microbiol. 11, 768108 (2021).

77. Timmerman, I., Daniel, A. E., Kroon, J. & van Buul, J. D. Leukocytes crossing the endothelium: A matter of communication. Int. Rev. Cell Mol. Biol. 322, 281– 329 (2016).

78. Shaftel, S. S., Griffin, W. S. T. & O’Banion, M. K. The role of interleukin-1 in neuroinflammation and Alzheimer disease: an evolving perspective. J. Neuroinflammation 5, 7 (2008).

79. Barnum, C. J. & Tansey, M. G. Modeling neuroinflammatory pathogenesis of Parkinson’s disease. Prog. Brain Res. 184, 113–132 (2010).

80. Gate, D. et al. CD4+ T cells contribute to neurodegeneration in Lewy body dementia. Science 374, 868–874 (2021).

81. van Langelaar, J., Rijvers, L., Smolders, J. & van Luijn, M. M. B and T cells driving multiple sclerosis: identity, mechanisms and potential triggers. Front. Immunol. 11, 760 (2020).

