## Supplementary material for "Microfluidics-free single-cell genomics reveals complex central-peripheral immune crosstalk in the mouse brain during peripheral inflammation": Table 1: percoll-vs-abdk_anova_table.html

|  | | Genotype effect | | Protocol effect | | Interaction | |
| --- | --- | --- | --- | --- | --- | --- | --- |
|  |  | *F*[1, 6] | *p* | *F*[1, 6] | *p* | *F*[1, 6] | *p* |
| Live cells | Raw count | 3.387 | 0.115 | 36.232 | 0.001 | 9.039 | 0.024 |
|  | % of live cells | 11.064 | 0.016 | 0.161 | 0.702 | 0.234 | 0.646 |
| CD45- | Raw count | 0.781 | 0.411 | 0.223 | 0.653 | 0.198 | 0.672 |
|  | % of live cells | 0.547 | 0.487 | 962.306 | < 0.001 | 33.516 | 0.001 |
| Microglia | Raw count | 9.930 | 0.02 | 81.385 | < 0.001 | 12.589 | 0.012 |
|  | % of live cells | 6.052 | 0.049 | 118.337 | < 0.001 | 0.892 | 0.381 |
| Dendritic cells | Raw count | 1.690 | 0.241 | 14.219 | 0.009 | 1.769 | 0.232 |
|  | % of live cells | 0.070 | 0.801 | 18.916 | 0.005 | 0.076 | 0.791 |
| Monocytes/macrophages | Raw count | 3.434 | 0.113 | 32.274 | 0.001 | 6.139 | 0.048 |
|  | % of live cells | 0.009 | 0.927 | 126.680 | < 0.001 | 3.163 | 0.126 |
| T-cells | Raw count | 7.243 | 0.036 | 0.440 | 0.532 | 0.756 | 0.418 |
|  | % of live cells | 5.199 | 0.063 | 1.761 | 0.233 | 1.692 | 0.241 |
| B-cells | Raw count | 0.734 | 0.425 | 4.441 | 0.08 | 0.868 | 0.388 |
|  | % of live cells | 0.068 | 0.802 | 3.544 | 0.109 | 0.208 | 0.664 |
| Neutrophils | Raw count | 0.242 | 0.641 | 0.768 | 0.414 | 0.136 | 0.725 |
|  | % of live cells | 1.903 | 0.217 | 1.902 | 0.217 | 0.779 | 0.411 |
