## Supplementary material for "Microfluidics-free single-cell genomics reveals complex central-peripheral immune crosstalk in the mouse brain during peripheral inflammation": Table 2: cross_entropy_results_collapseLPS.html

|  | Kullback-Leibler divergence | Holm-adjusted *p*-value |
| --- | --- | --- |
| B-lymphocytes | 0.056 | 0.23 |
| CD45- cells | 0.039 | 0.064 |
| Dendritic cells | 0.045 | 0.497 |
| Granulocytes | 0.132 | < 0.001 |
| Macrophages | 0.092 | < 0.001 |
| Microglia | 0.107 | < 0.001 |
| Monocytes | 0.046 | 0.007 |
| Natural killer cells | 0.052 | 0.941 |
| T-cells | 0.029 | 0.993 |
