## Supplementary material for "Microfluidics-free single-cell genomics reveals complex central-peripheral immune crosstalk in the mouse brain during peripheral inflammation": Table 3: microbead_dilution_anova_table.html

|  | | Effect of CD45 antibody clone | | | Effect of MicroBead dilution | | | Interaction | | |
| --- | --- | --- | --- | --- | --- | --- | --- | --- | --- | --- |
|  |  | df  (numerator, denominator) | *F* | *p* | df  (numerator, denominator) | *F* | *p* | df  (numerator, denominator) | *F* | *p* |
| Live cell count | Positive | 1.199, 10.793 | 31.725 | < 0.001 | 2, 9 | 25.336 | < 0.001 | 2.398, 10.793 | 7.069 | 0.009 |
|  | Negative | 2, 18 | 0.504 | 0.612 | 2, 9 | 0.885 | 0.446 | 4, 18 | 0.958 | 0.454 |
|  | Non-Enriched | 2, 6 | 2.335 | 0.178 | — | — | — | — | — | — |
| Microglial proportion | Positive | 2, 18 | 3.943 | 0.038 | 2, 9 | 0.289 | 0.755 | 4, 18 | 1.116 | 0.38 |
|  | Negative | 2, 18 | 2.499 | 0.11 | 2, 9 | 1.475 | 0.279 | 4, 18 | 1.342 | 0.293 |
|  | Non-Enriched | 2, 6 | 6.161 | 0.035 | — | — | — | — | — | — |
| Non-microglia myeloid proportion | Positive | 1.225, 11.024 | 128.48 | < 0.001 | 2, 9 | 20.705 | < 0.001 | 2.45, 11.024 | 8.047 | 0.005 |
|  | Negative | 2, 18 | 13.849 | < 0.001 | 2, 9 | 1.989 | 0.193 | 4, 18 | 0.438 | 0.78 |
|  | Non-Enriched | 2, 6 | 26.251 | 0.001 | — | — | — | — | — | — |
| Lymphocyte proportion | Positive | 2, 18 | 80.506 | < 0.001 | 2, 9 | 11.408 | 0.003 | 4, 18 | 2.245 | 0.105 |
|  | Negative | 1.15, 10.348 | 93.168 | < 0.001 | 2, 9 | 0.971 | 0.415 | 2.3, 10.348 | 1.867 | 0.201 |
|  | Non-Enriched | 2, 6 | 43.285 | < 0.001 | — | — | — | — | — | — |
| CD45- cell proportion | Positive | 2, 18 | 13.573 | < 0.001 | 2, 9 | 1.727 | 0.232 | 4, 18 | 1.818 | 0.169 |
|  | Negative | 2, 18 | 15.8 | < 0.001 | 2, 9 | 0.917 | 0.434 | 4, 18 | 3.2 | 0.038 |
|  | Non-Enriched | 2, 6 | 17.519 | 0.003 | — | — | — | — | — | — |
