## Supplementary figures and images for "Microfluidics-free single-cell genomics reveals complex central-peripheral immune crosstalk in the mouse brain during peripheral inflammation"

### Figure S1

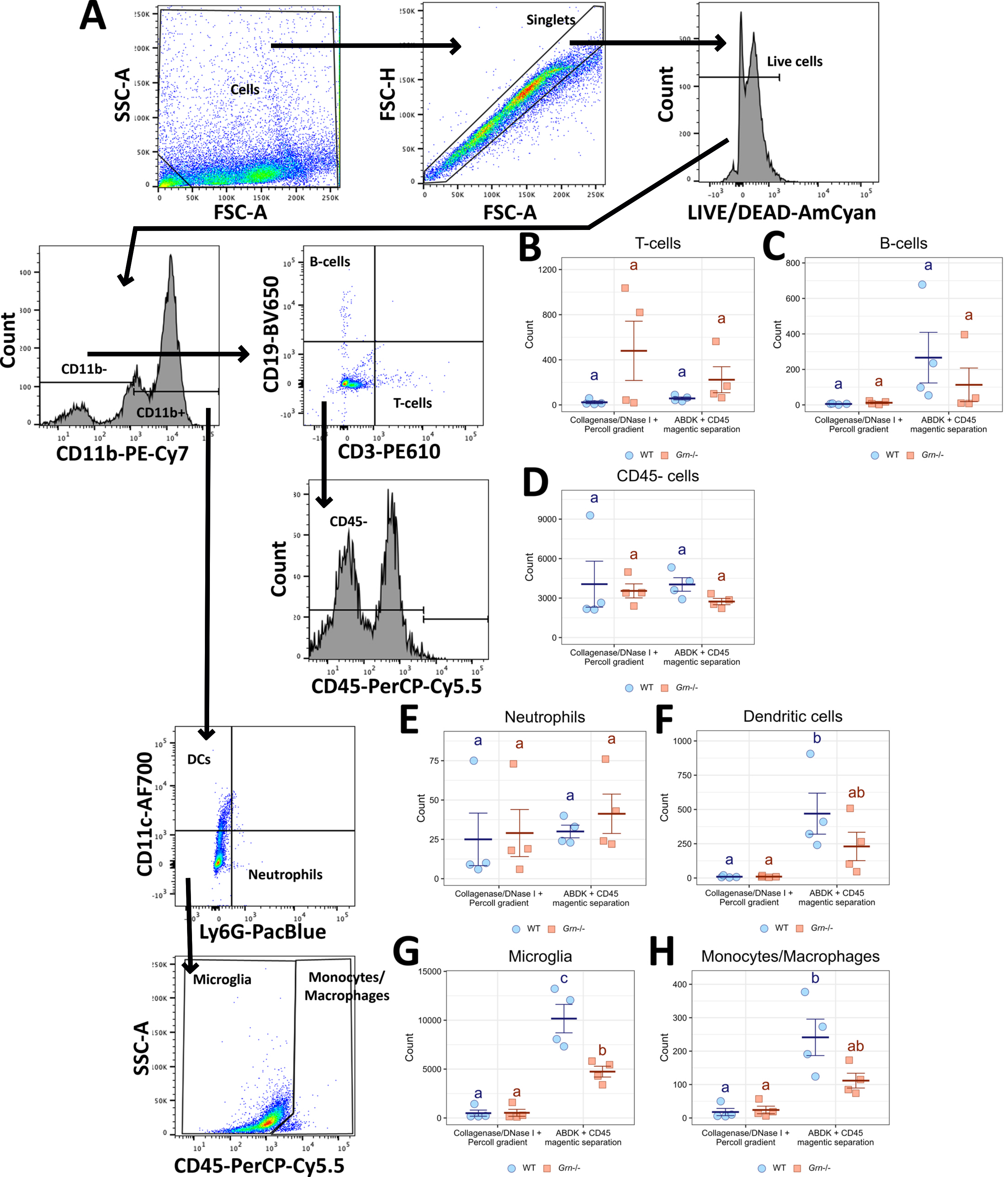

### Figure S2

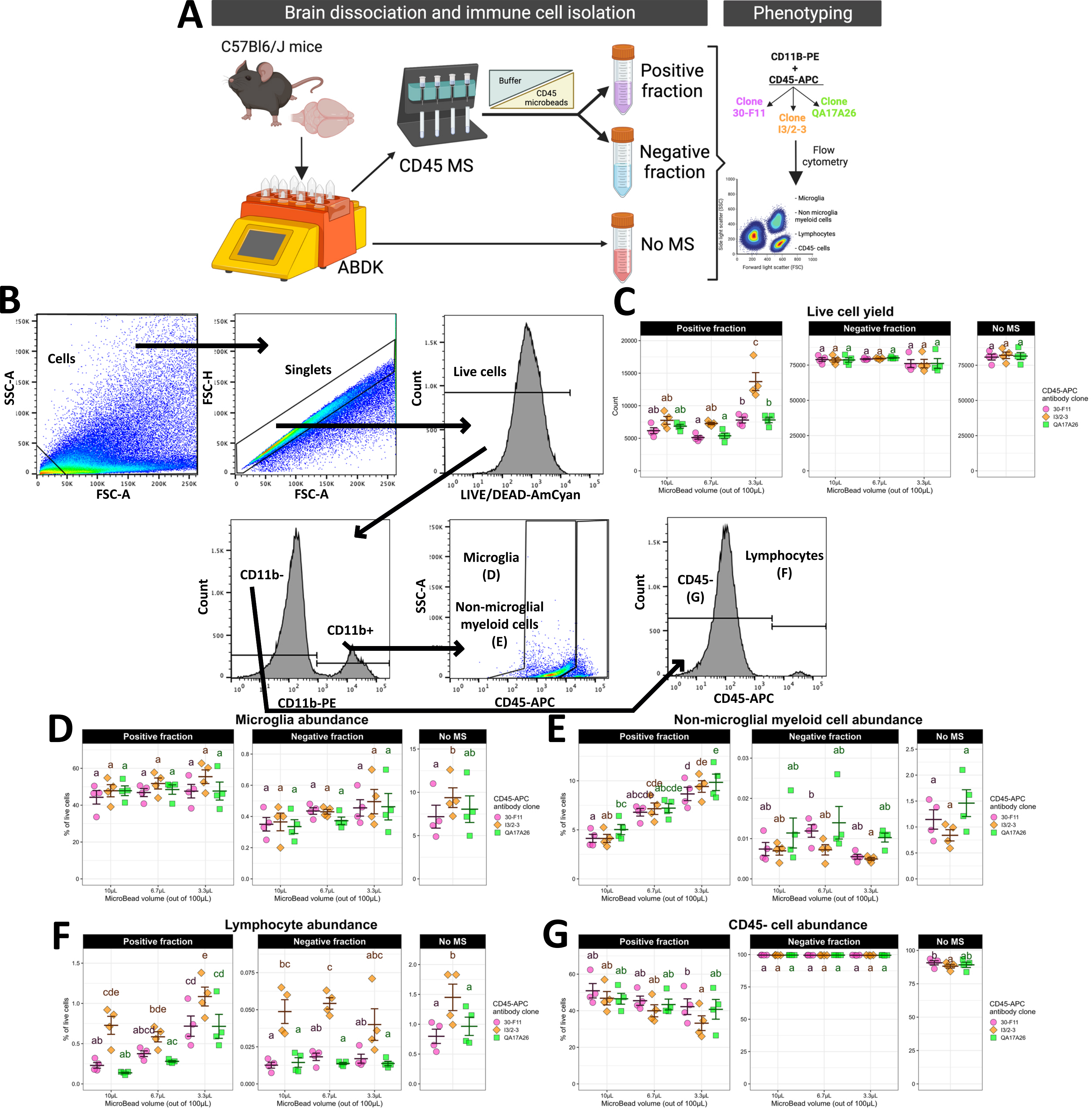

### Figure S3

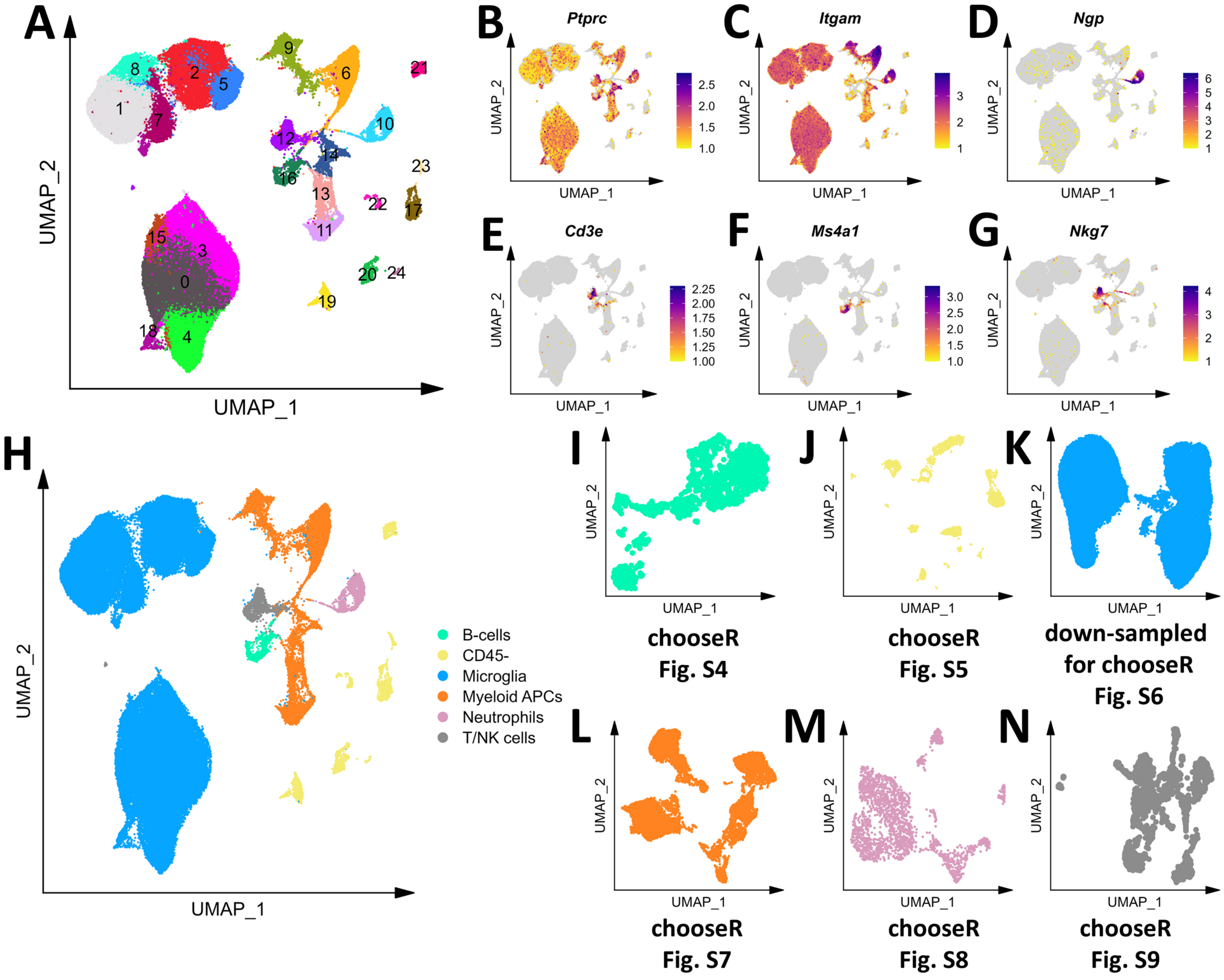

### Figure S4

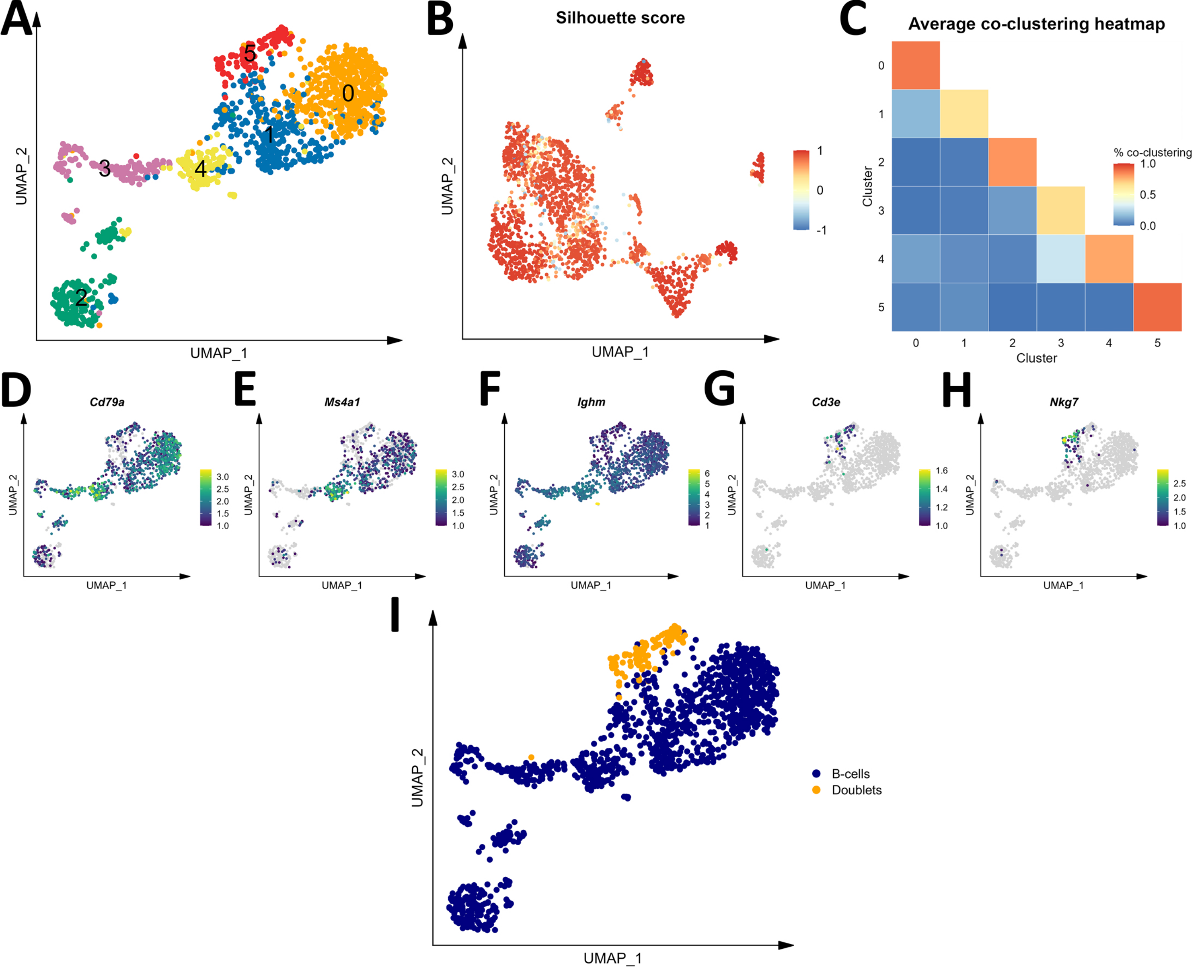

### Figure S5

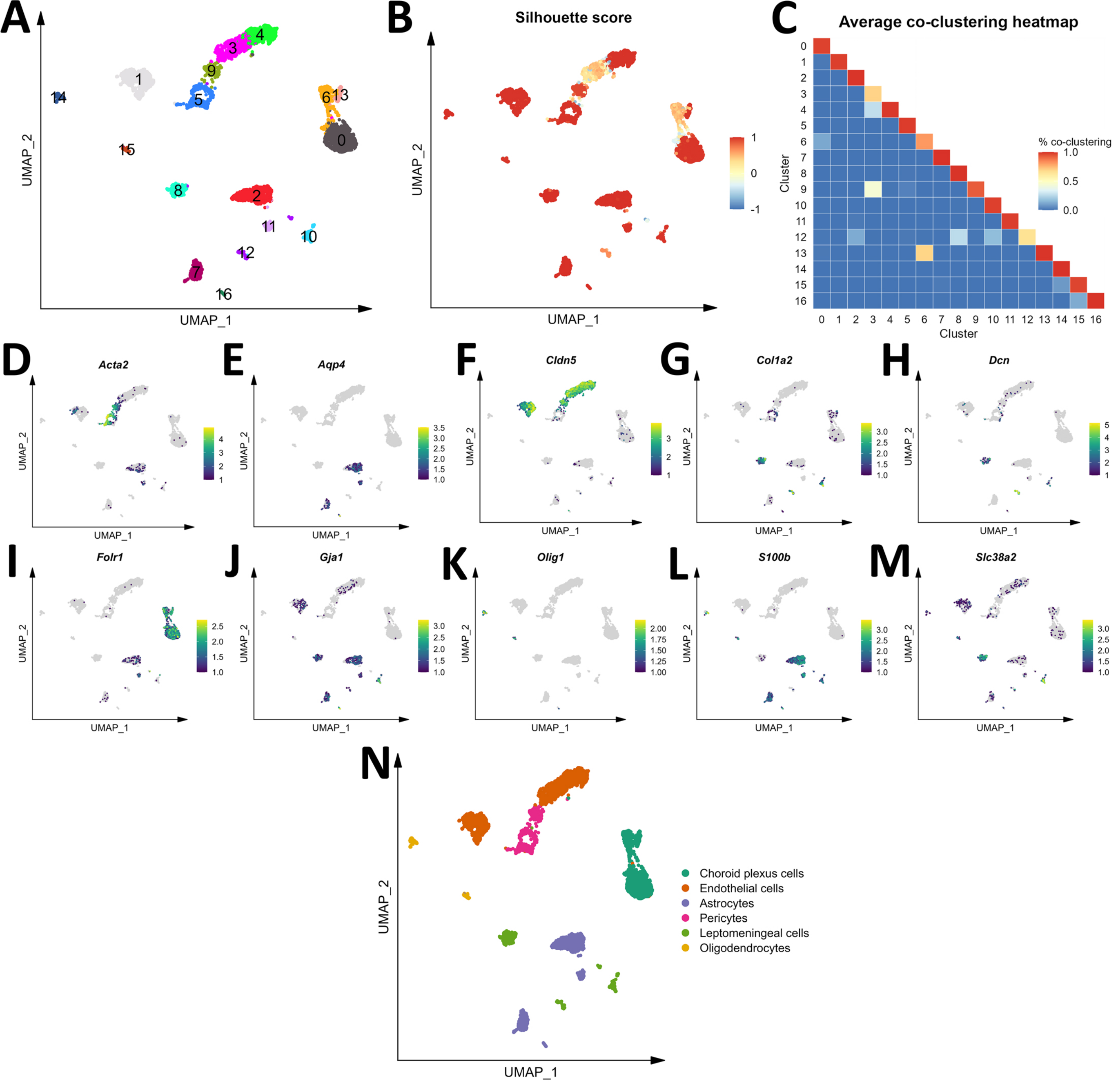

### Figure S6

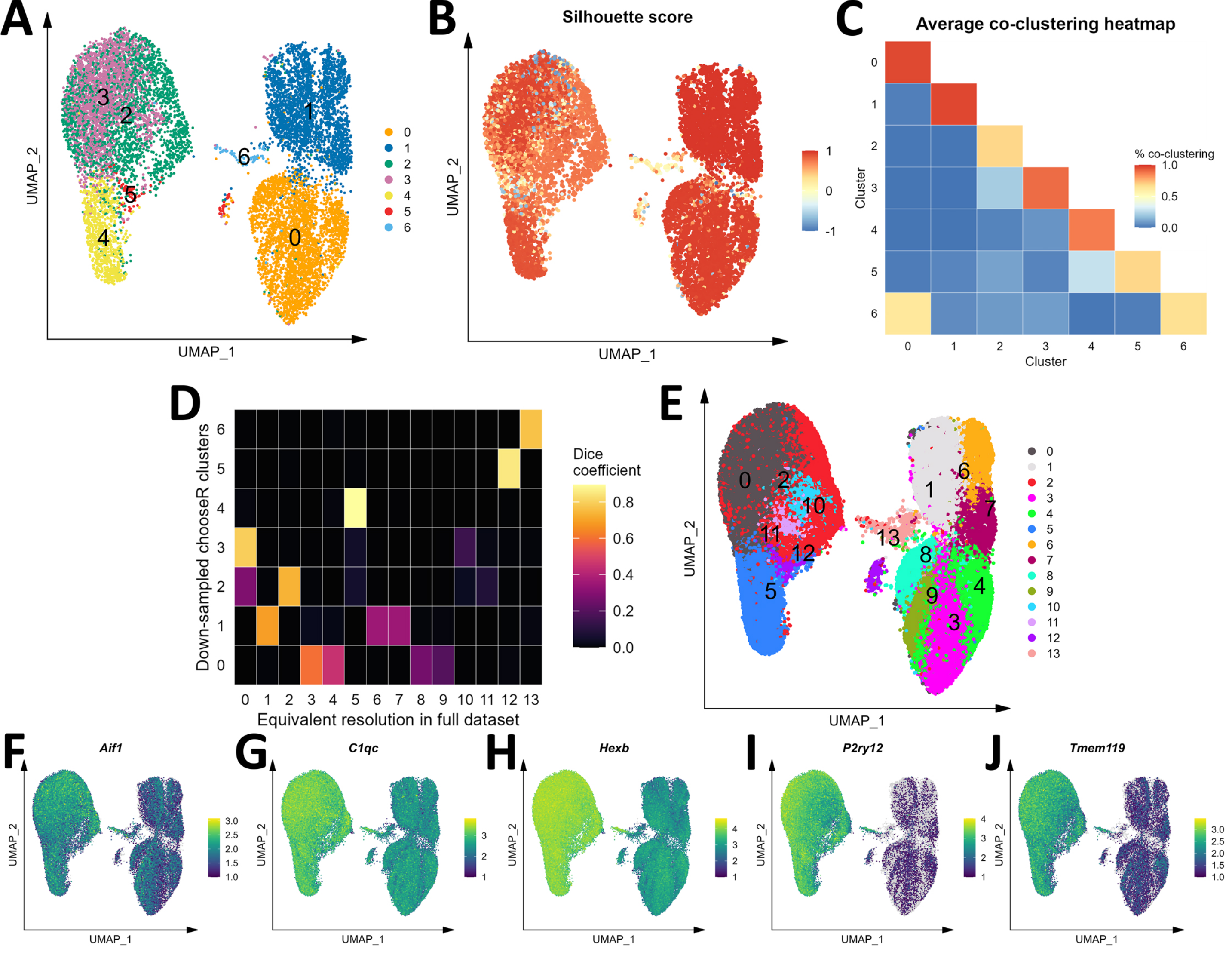

### Figure S7

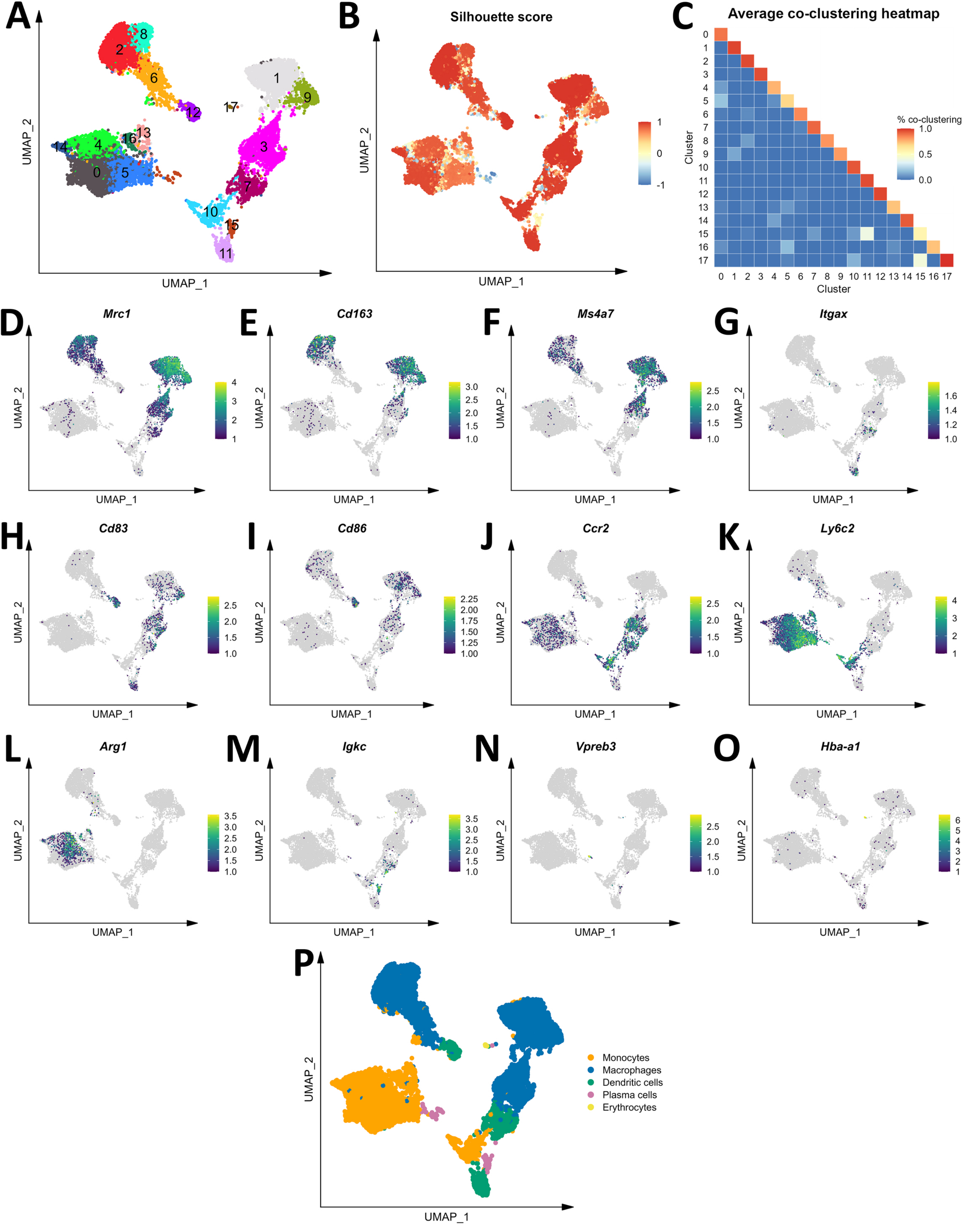

### Figure S8

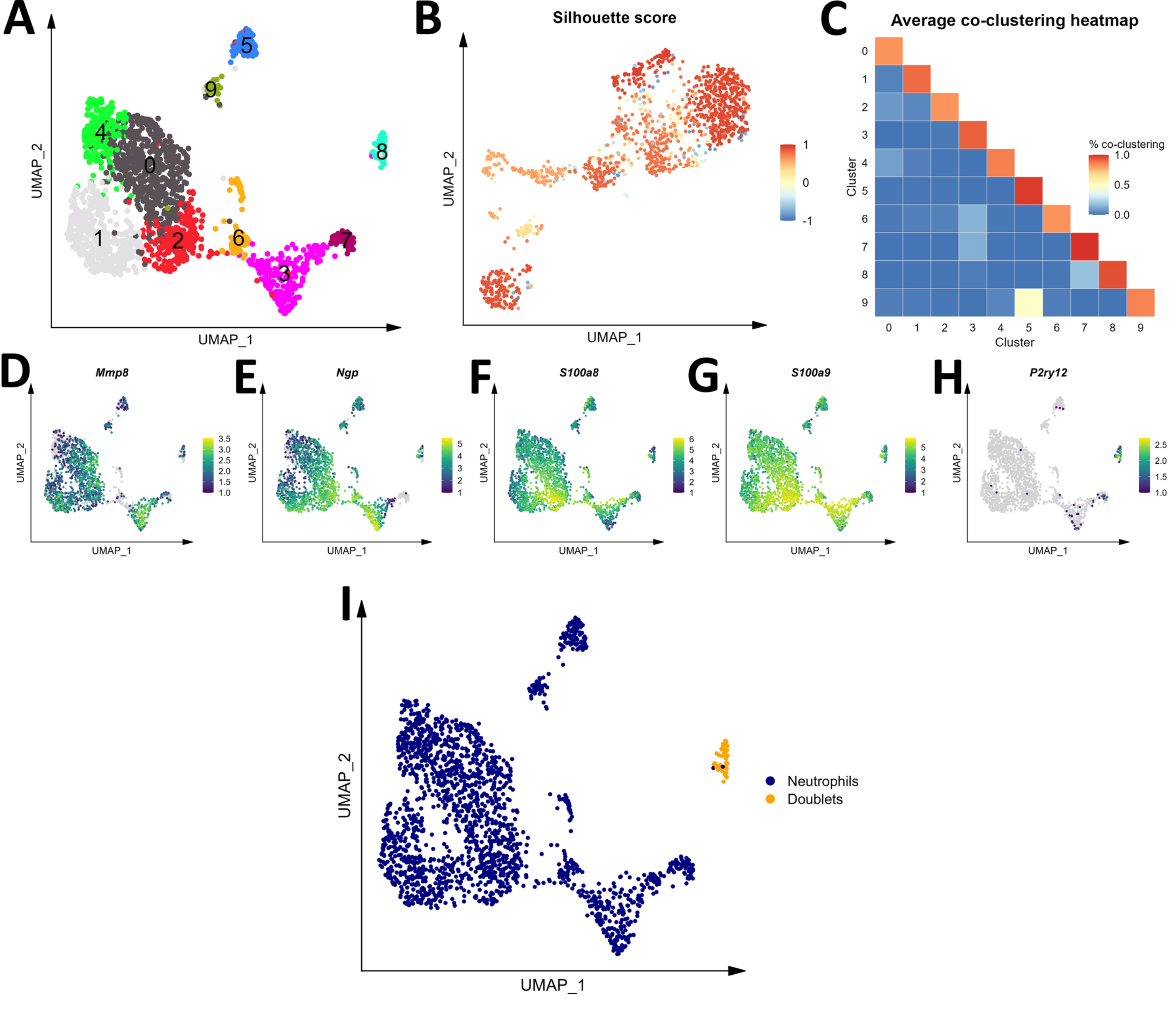

### Figure S9

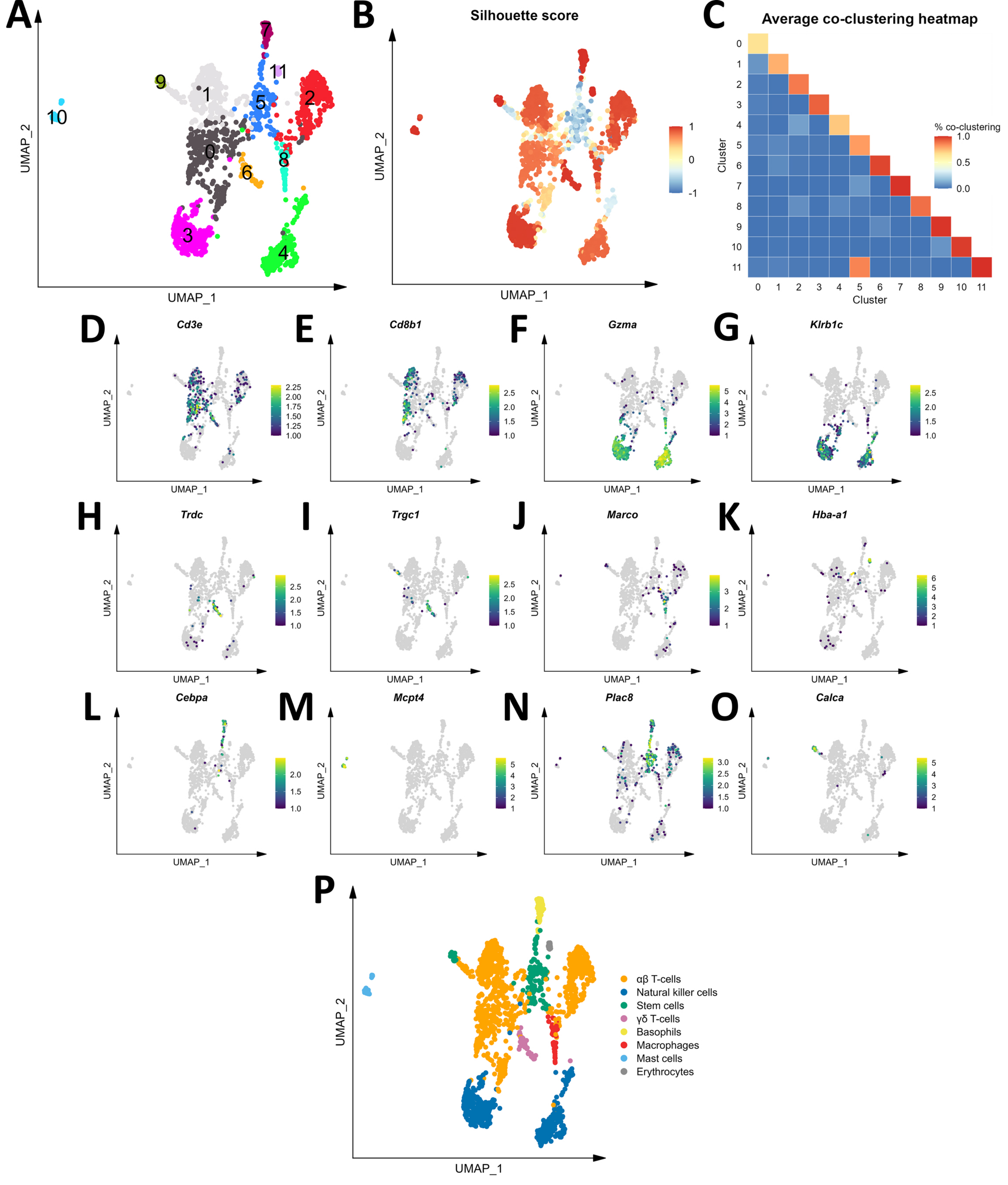

### Figure S10

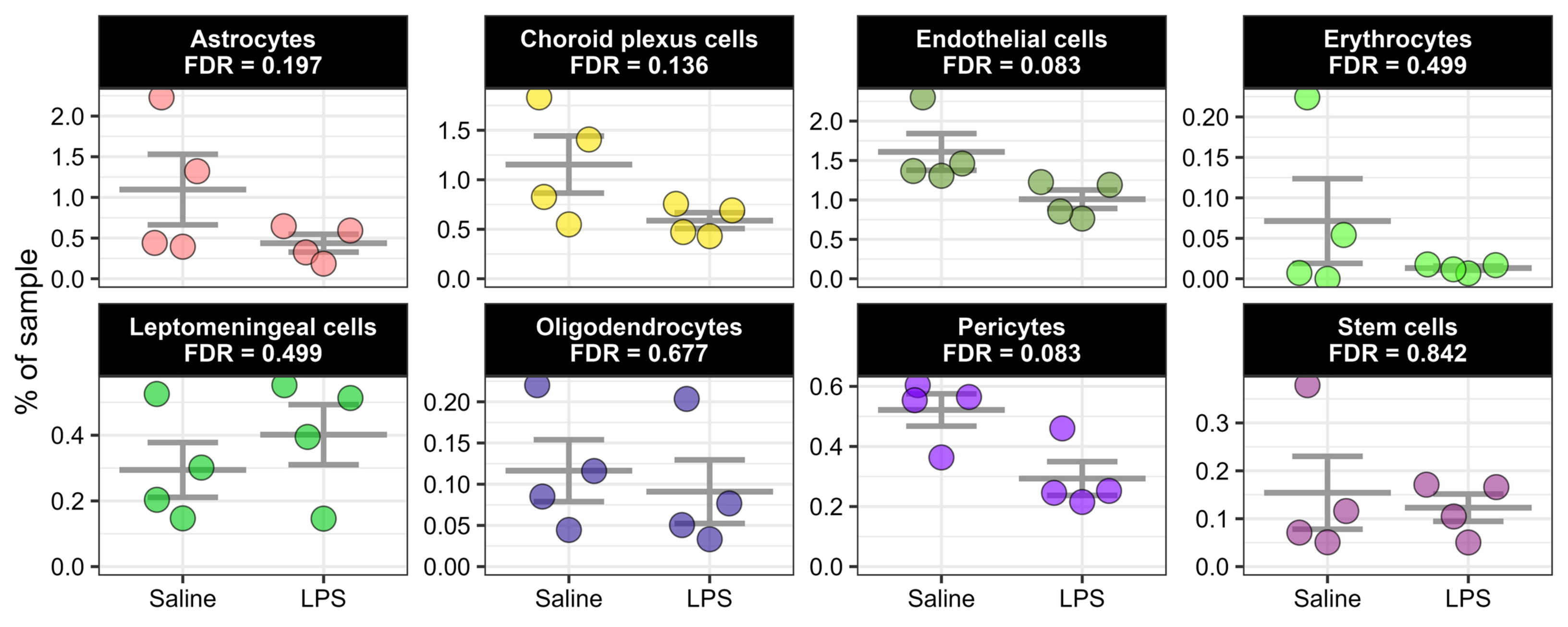

### Figure S11

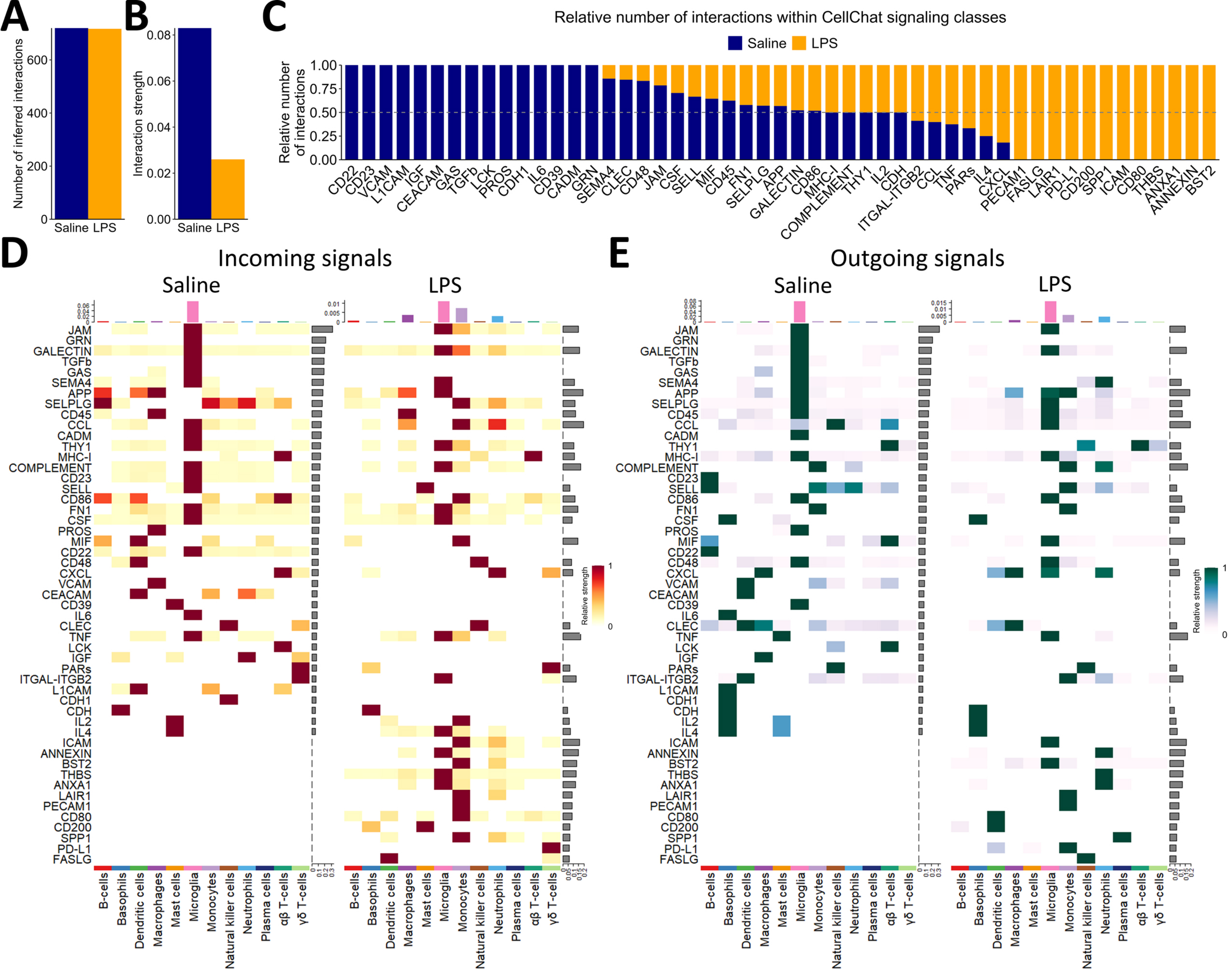

### Figure S12

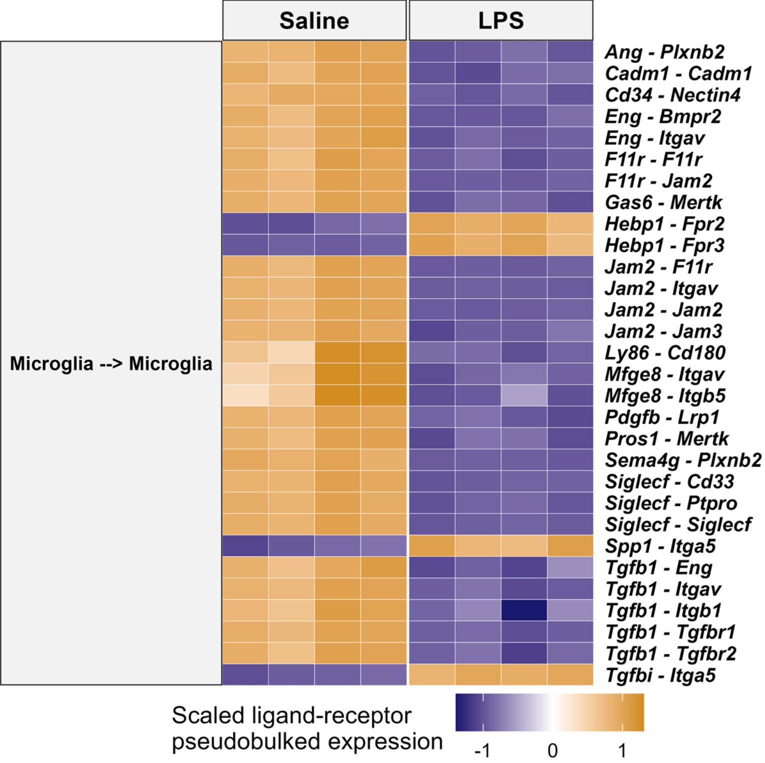
